# A 9-kDa matricellular SPARC fragment released by cathepsin D exhibits pro-tumor activity in the triple-negative breast cancer microenvironment

**DOI:** 10.1101/2020.10.22.350082

**Authors:** Lindsay B Alcaraz, Aude Mallavialle, Timothée David, Danielle Derocq, Frédéric Delolme, Cindy Dieryckx, Florence Boissière-Michot, Joëlle Simony-Lafontaine, Stanislas Du Manoir, Pitter F. Huesgen, Christopher M. Overall, Sophie Tartare-Deckert, William Jacot, Thierry Chardès, Séverine Guiu, Pascal Roger, Thomas Reinheckel, Catherine Moali, Emmanuelle Liaudet-Coopman

**Author notes:** Corresponding author: E Liaudet-Coopman.

## Abstract

Extracellular matrix (ECM) remodeling by proteases results in the release of protein fragments that promote tumor progression and metastasis. The protease cathepsin D (cath-D), a marker of poor prognosis in triple-negative breast cancer (TNBC), is aberrantly secreted in the tumor microenvironment. Using degradomic analyses by TAILS, we discovered that the matricellular protein SPARC is a substrate of extracellular cath-D. *In vitro*, cath-D induced limited proteolysis of SPARC C-terminal extracellular Ca^2+^ binding domain at acidic pH, leading to the production of SPARC fragments (34-, 27-, 16-, 9-, and 6-kDa). Similarly, cath-D secreted by TNBC cells cleaved fibroblast- and cancer cell-derived SPARC at the tumor pericellular acidic pH. SPARC cleavage also occurred in TNBC tumors. Among these fragments, only the 9-kDa SPARC fragment inhibited TNBC cell adhesion and spreading on fibronectin, and stimulated their migration, endothelial transmigration, and invasion. Our study establishes a novel crosstalk between proteases and matricellular proteins in the tumor microenvironment through limited proteolysis of SPARC, revealing a novel targetable 9-kDa bioactive SPARC fragment for new TNBC treatments.

## INTRODUCTION

Breast cancer (BC) is one of the leading causes of death in women in developed countries. Triple-negative breast cancer (TNBC), defined by the absence of estrogen receptor (ER), progesterone receptor (PR) and human epidermal growth factor receptor 2 (HER-2) overexpression and/or amplification, accounts for 15-20% of all BC cases [1]. Chemotherapy is the primary systemic treatment, but resistance to this treatment is common [1]. Thus, tumor-specific molecular targets are urgently needed to develop alternative therapeutic strategies for TNBC.

Tumor progression has been recognized as the product of the dynamic crosstalk between tumor cells and the surrounding tumor stroma [2]. Cancer cells interact dynamically with several cell types within the extracellular matrix (ECM), such as fibroblasts, endothelial cells, adipocytes and infiltrating immune cells. Stromal and tumor cells exchange ECM proteins and bioactive fragments, enzymes, growth factors, and cytokines that modify the local ECM, thus affecting cell-matrix and cell-cell adhesion, and promoting migration and invasion, proliferation and survival of stromal and tumor cells [3, 4]. In the last decade, it has become increasingly evident that cancer cells create a pericellular microenvironment where several protease families, such as metalloproteinases, serine proteases, cysteine and aspartic cathepsins cooperate to form a pro-tumorigenic proteolytic network [5]. These proteases participate in the remodeling of the surrounding ECM during tumor progression and metastasis formation, releasing a number of bioactive ECM fragments that can support carcinogenesis [3, 4]. Therefore, the identification of protease repertoires by Omics strategies, such as N-Terminal Amine Isotopic Labeling of Substrates (TAILS)-based degradome analysis [6], is important for the discovery of novel targetable bioactive ECM protein fragments in cancer.

Human cathepsin D (cath-D) is a ubiquitous, lysosomal, aspartic endoproteinase that is proteolytically active at acidic pH. Cath-D expression levels in BC [7–9] and TNBC [10, 11] correlate with poor prognosis. We recently reported that cath-D is a tumor-specific extracellular target in TNBC and is suitable for antibody-based therapy [12]. Cath-D over-production by BC and TNBC cells leads to hypersecretion of the 52-kDa cath-D precursor in the tumor microenvironment [11, 13]. Purified 52-kDa cath-D auto-activates in acidic conditions, giving rise to a catalytically active 51-kDa pseudo-cath-D form that retains the 18 residues (27-44) of the pro-segment [14]. Cath-D affects the tumor and its microenvironment by increasing BC cell proliferation [13, 15–18], and by stimulating mammary fibroblast outgrowth [19, 20], angiogenesis [15, 21], and metastasis formation [17]. However, little is known about the molecular mechanisms and the substrates involved in these processes.

In this study we investigated cath-D substrate repertoire in the TNBC tumor microenvironment by TAILS-based degradomic analysis, and identified the matricellular protein Secreted Protein Acidic and Rich in Cysteine (SPARC), also known as osteonectin or basement membrane 40 (BM40). SPARC is a Ca^2+^-binding glycoprotein that regulates ECM assembly and deposition, growth factor signaling, and ECM-cell interactions [22–25]. In cancer, SPARC is mainly secreted by the neighboring stroma, but also by cancer cells [26–28]. SPARC plays an oncogenic or a tumor-suppressive role in function of the cancer type [29, 30]. For instance, in BC, SPARC has a pro-tumorigenic role and has been associated with worse prognosis [27, 31–36]; however, other studies reported anti-tumorigenic functions [37–39]. SPARC includes three different structural and functional modules: the N-terminal acidic domain, followed by the follistatin-like domain, and the C-terminal extracellular Ca^2+^ binding domain [24]. In this study, we found that, in the TNBC acidic microenvironment, cath-D cleaved SPARC exclusively in its C-terminal extracellular Ca^2+^ binding domain, releasing five main fragments (34-, 27-, 16-, 9-, and 6-kDa). Among these fragments, the 9-kDa C-terminal SPARC fragment (amino acids 235-303) had greater oncogenic activity than full-length (FL) SPARC, highlighting the importance of limited proteolysis of matricellular proteins in the TNBC microenvironment. This knowledge might pave the way to the development of strategies to target bioactive matricellular proteins in TNBC.

## RESULTS

### SPARC is an extracellular protein cleaved in the TNBC microenvironment

To investigate the cath-D extracellular substrate repertoire in the TNBC microenvironment, we designed a coculture assay in which cath-D-secreting MDA-MB-231 TNBC cells are grown with human mammary fibroblasts (HMF) (Fig. 1A). To determine cath-D impact on extracellular protein processing, we analyzed by TAILS the secreted proteome in supernatants of MDA-MB-231 cells and HMFs co-cultures after incubation at acidic pH in the presence or absence of pepstatin A, an aspartic protease inhibitor. We identified 4130 unique N-terminal peptides, among which 3091 could be quantified (Fig. 1B), from 1582 unique proteins. A 2-fold change of the without pepstatin A/with pepstatin A ratio after 60 min of incubation was used as cutoff to determine N termini substantially affected at acidic pH (Supplementary Fig. 1). Based on this criterion, 200 neo-N termini were more abundant in the absence of pepstatin A, while the abundance of 5 mature protein N-termini was reduced in the absence of pepstatin (Supplementary Table 1). Several SPARC peptides were among the peptides showing the most prominent changes in abundance between conditions (Fig. 1B, Table 1 and Supplementary Table 1). To determine whether SPARC was a putative cath-D substrate, we first confirmed by western blotting of the same samples the reduction of FL SPARC level and the presence of new fragments with lower molecular weight in the absence of pepstatin A (Fig. 1C). These data suggested that SPARC is cleaved in the TNBC extracellular environment.

**Figure 1.**
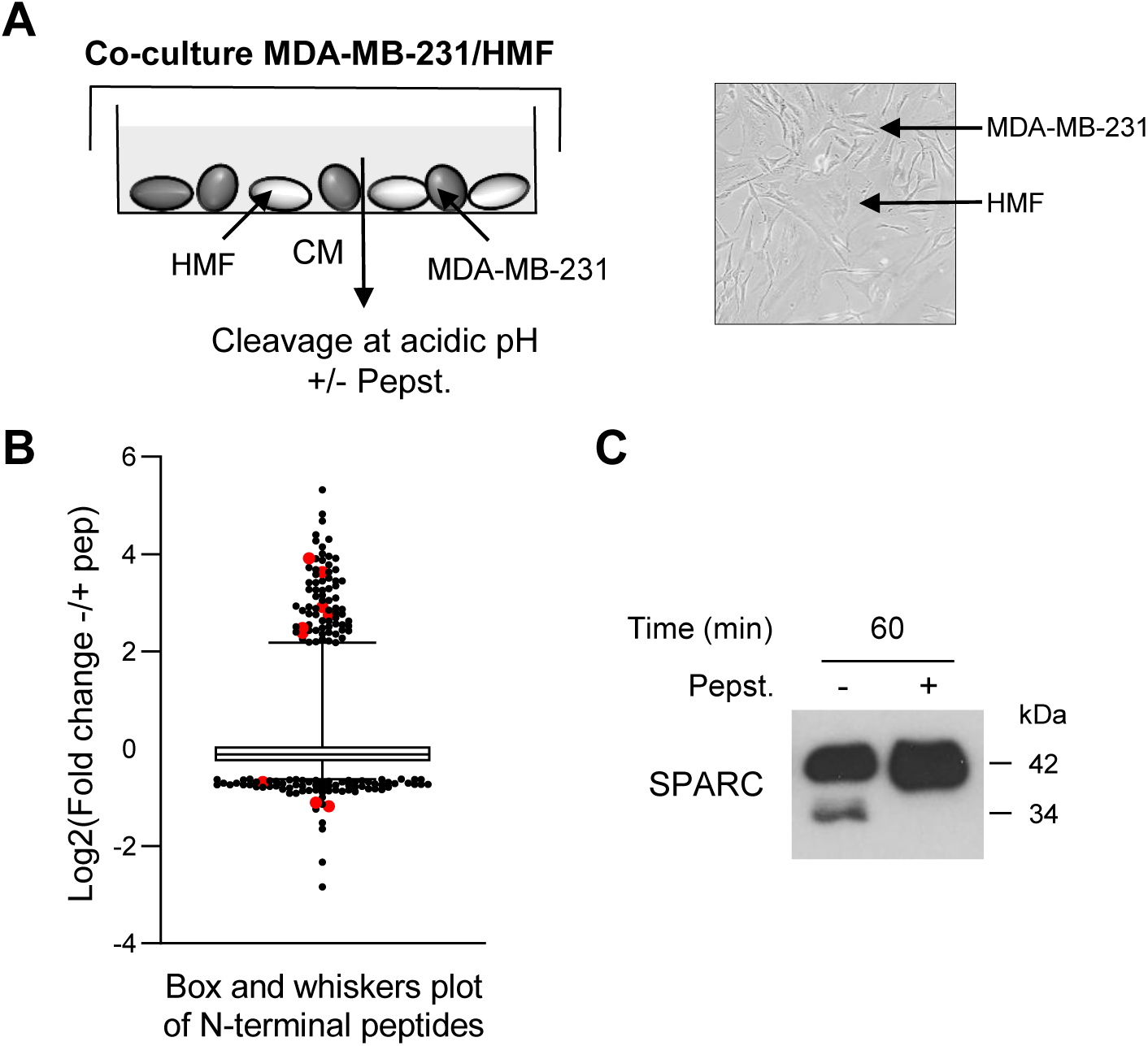
Identification of SPARC as an extracellular protein cleaved in the TNBC microenvironment at acidic pH. **(A) Experimental set-up of the MDA-MB-231 cell-HMF co-culture system.** MDA-MB-231 TNBC cells and HMFs were co-cultured in serum-free DMEM without sodium bicarbonate and phenol red and buffered with 50 mM HEPES [pH 7.5] at 37°C for 24h. The 24h-conditioned medium was then concentrated to 0.2 mg/ml, and incubated in cleavage buffer with or without pepstatin A (Pepst.) (12.5 µM) at pH 5.5 and 37° for 60 min. A representative image of an MDA-MB-231/HMF co-culture is shown in the right panel (x 200). **(B) Box and whisker plot of the normalized ratios of the N-terminal peptides identified by TAILS in the MDA-MB-231/HMF co-culture secretome.** 3091 peptides quantified at t = 60 min of incubation with cleavage buffer were used to generate the graph. The without/with pepstatin A ratios corresponding to SPARC peptides are highlighted in red; whiskers correspond to the 2.5^th^ and 97.5^th^ percentiles. **(C) Validation of SPARC cleavage in the MDA-MB-231/HMF co-culture secretome.** Secretome samples (2 µg) from the MDA-MB-231/HMF co-culture incubated in cleavage buffer with or without pepstatin A (Pepst.) (12.5 µM) at pH 5.5 and at 37° for 60 min were separated on 13.5% SDS-PAGE followed by immunoblotting with anti-SPARC antibody (clone AON-5031).

**Table 1.** Sequences of the SPARC fragments identified by TAILS and ATOMS. High-confidence peptides with N-terminal iTRAQ labelling matching the FL SPARC protein (Uniprot accession number P09486) identified by TAILS in the conditioned medium of co-cultured MDA-MB-231/HMFs, or by iTRAQ-ATOMS after *in vitro* cleavage of recombinant SPARC by recombinant cath-D. Peptides defining cleavage sites with iTRAQ ratios >2 and <0.5 for TAILS and ATOMS, respectively, are shown. CM, conditioned medium; pepst., pepstatin A; *, according to the silver staining.

|  |  |  | TAILS<br>(SPARC cleavages<br>from conditioned<br>media of<br>cocultures) | ATOMS<br>(*in vitro* cleavages of recombinant<br>SPARC by recombinant cath-D) |  |
| --- | --- | --- | --- | --- | --- |
| SPARC peptides | SPARC<br>fragments* | Enzyme<br>used for<br>sample<br>digestion | (CM -pepst./CM<br>+pepst.) ratio<br>(number of<br>peptide<br>identifications) | (Pseudo-cathD +<br>SPARC<br>- pepst.<br>/Pseudo-cath-D<br>+ SPARC +pepst)<br>ratio (number<br>of peptide<br>identifications) | (Mature cath-D +<br>SPARC<br>-pepst.<br>/Mature cath-D +<br>SPARC + pepst.)<br>ratio (number of<br>peptide<br>identifications) |
| LDSELTEFPLR [156-166] |  | Trypsin | 0.4 (6) |  |  |
| DWLKNVLVTLYER [169-181] |  | Trypsin | 0.5 (3) |  |  |
| VTLYERDEDNNLLTEK [176-191] | 16-kDa | Trypsin |  | 12.1 (5) | 14.3 (5) |
| YERDEDNNLLTEK [179-191] |  | Trypsin |  | 13.7 (16) | 11.8 (16) |
| YERDEDNNLLTEKQK [179-193] |  | Trypsin |  | 21.1 (15) | 16.8 (15) |
| EAGDHPVELLAR [207-218] | 27-kDa | Trypsin | 3.4 (2) | 11.3 (6) | 19.3 (6) |
| PVHWQFGQLDQHPIDGYLSHTELA PLR [230-256] | 9-kDa | Trypsin | 5.6 (8) |  |  |
| WQFGQLDQHPIDGYLSHTELA PLR [233-256] |  | Trypsin | 3.1 (3) |  |  |
| FGQLDQHPIDGYLSHTELA PLR [235-256] |  | Trypsin |  | 10.6 (5) | 5.9 (5) |
| GQLDQHPIDGYLSHTELA PLR [236-256] |  | Trypsin | 12.4 (15) | 2 (16) | 2.3 (16) |
| QLDQHPIDGYLSHTELA PLR [237-256] |  | Trypsin | 15.1 (5) | 3.5 (5) | 2.2 (5) |
| DQHPIDGYLSHTELA PLR [239-256] |  | Trypsin | 7.6 (18) | 8.4 (18) | 8.1 (18) |
| YLSHTELA PLR [246-256] |  | Trypsin | 4.4 (4) | 18.8 (8) | 5.7 (8) |
| LSHTELA PLR [247-256] |  | Trypsin | 2.4 (1) |  |  |
| SHTELA PLR [248-256] |  | Trypsin |  | 2.7 (5) | 1.6 (5) |
| PLIPMEHCTTR [258-268] | 6- and 34-kDa | Trypsin | 5.9 (3) | 22.5 (15) | 4.3 (15) |
| APQQEALPDE [18-27] |  | Glu-C |  |  | 0.98 (2) |
| APQQEALPDETE [18-29] |  | Glu-C |  |  | 0.88 (13) |
| APQQEALPDETEVVE [18-32] |  | Glu-C |  |  | 1.12 (6) |
| APQQEALPDETEVVEE [18-33] |  | Glu-C |  |  | 1.1 (140) |
| APQQEALPDETEVVEETVAE [18-37] |  | Glu-C |  |  | 1.14 (18) |
| VTLYERDEDNNLLTE [176-190] | 16.4-kDa | Glu-C |  |  | **14.3** (12) |
| YERDEDNNLLTE [179-190] |  | Glu-C |  |  | **13.1** (43) |
| FGQLDQHPIDGYLSHTE [235-251] | 9-kDa | Glu-C |  |  | **9.2** (3) |
| GQLDQHPIDGYLSHTE [236-251] |  | Glu-C |  |  | **10.4** (12) |
| DQHPIDGYLSHTE [239-251] |  | Glu-C |  |  | 4.7 (6) |

Ratio; <0.5 or >2 for trypsin ; Ratio; <0.5 or >2 in bold for Glu-C; CM, conditioned medium ; pepst., pepstatin A ; \*, according to silver staining.

### *In vitro,* cath-D cleaves SPARC extracellular Ca^2+^ binding domain at acidic pH

We next investigated whether recombinant cath-D can cleave recombinant SPARC *in vitro* at acidic pH. At pH 5.9, SPARC was cleaved by catalytically active 51-kDa pseudo-cath-D in a time-dependent manner (Fig. 2A). Moreover, experiments in which pH was gradually reduced from 6.8 to 5.5 showed progressive limited proteolysis of SPARC at lower pH (Fig. 2B). In these two experiments, pepstatin A, inhibited SPARC cleavage by cath-D (Fig. 2A-B). By amino-terminal oriented mass spectrometry of substrates (ATOMS) analysis, we found that at pH 5.9, SPARC was cleaved by the 51-kDa cath-D form exclusively in its extracellular Ca^2+^ binding domain, releasing five main SPARC fragments (34-, 27-, 16-, 9-, and 6-kDa) that could be detected by silver staining (Fig. 2C-E, Table 1). We detected SPARC cleavage fragments of similar size also after incubation with the fully mature 34+14-kDa cath-D form at pH 5.9 (Fig. 2C-E, Table 1). Thus, *in vitro,* cath-D triggers SPARC limited proteolysis exclusively in its extracellular Ca^2+^ binding domain in an acidic environment.

**Figure 2.**
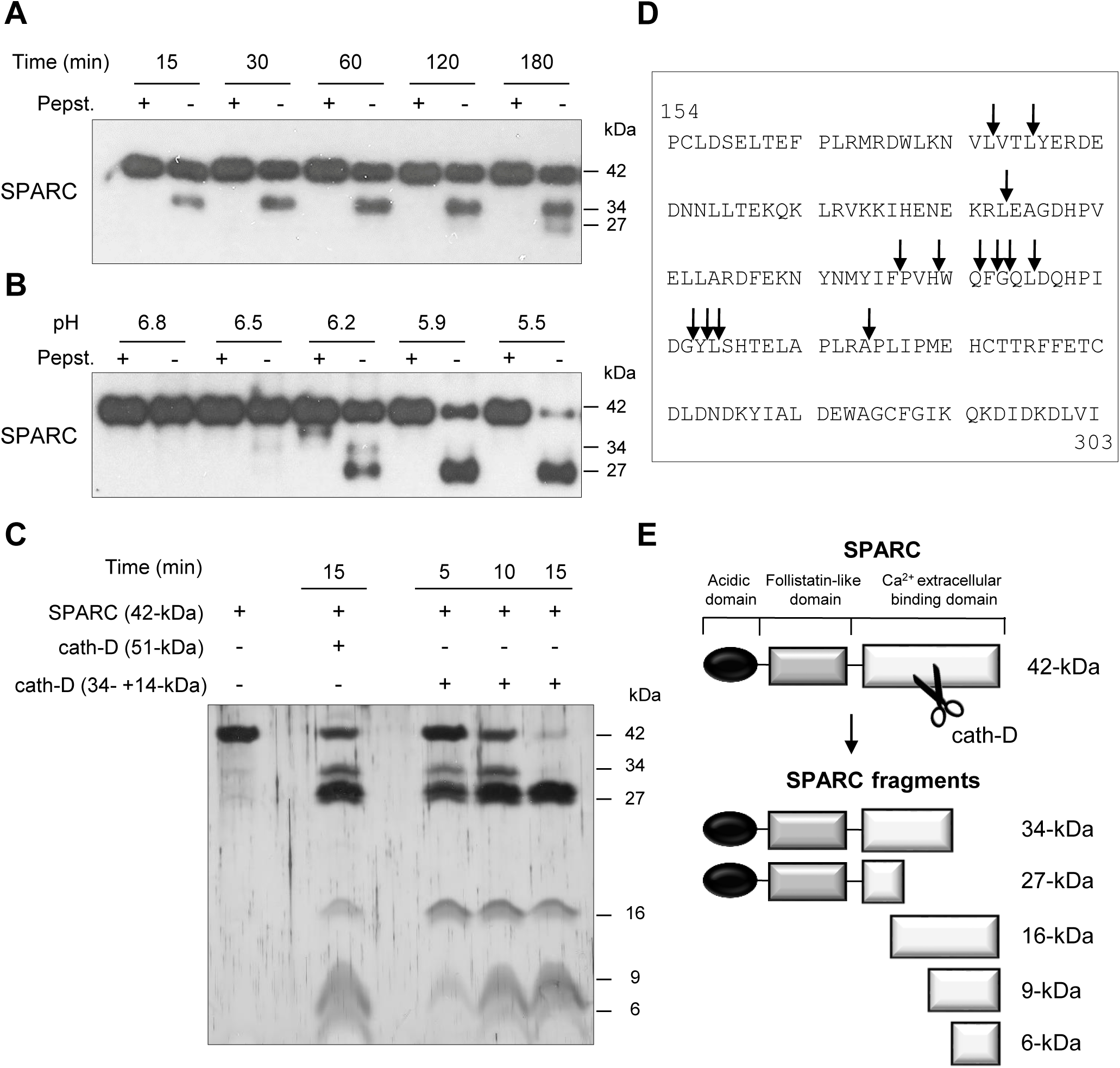
Cleavage of the extracellular Ca^2+^ binding domain of human SPARC by human cath-D at acidic pH. **(A) Time-course of cath-D-induced SPARC cleavage.** Recombinant human FL SPARC was incubated with recombinant human auto-activated pseudo-cath-D (51-kDa) in cleavage buffer at pH 5.9 with or without pepstatin A (Pepst.) at 37°C for the indicated times. SPARC cleavage was analyzed by 13.5% SDS-PAGE and immunoblotting with an anti-SPARC antibody (clone AON-5031). **(B) pH dependence of cath-D-induced SPARC cleavage.** Recombinant human FL SPARC was incubated with recombinant human auto-activated pseudo-cath-D (51-kDa) in cleavage buffer with or without pepstatin A (Pepst.) at the indicated pH at 37°C overnight. SPARC cleavage was analyzed as in (**A**). **(C) Detection of the cath-D-induced SPARC fragments by silver staining.** Recombinant SPARC was incubated with recombinant auto-activated pseudo-cath-D (51-kDa) or fully-mature cath-D (34+14-kDa) at pH 5.9 for the indicated times. SPARC cleavage was analyzed by 17% SDS-PAGE and silver staining. **(D) Cath-D cleavage sites in SPARC extracellular Ca^2+^ binding domain.** The entire C-terminal extracellular Ca^2+^ binding domain of human SPARC (amino acids 154-303) is shown. SPARC cleaved peptides generated in the extracellular Ca^2+^ binding domain by auto-activated pseudo-cath-D (51-kDa) and fully-mature (34+14-kDa) cath-D at pH 5.9 were resolved by iTRAQ-ATOMS. Arrows, cleavage sites. **(E) Schematic representation of the SPARC fragments generated by cath-D according to (C) and (D).**

### SPARC and cath-D expression in TNBC

To study the pathophysiological relevance of the SPARC/cath-D interplay in TNBC, we first assessed *SPARC* and *CTSD* (the gene encoding cath-D) expression in TNBC samples from 255 patients using an online survival analysis [40]. High *CTSD* mRNA level was significantly associated with shorter recurrence-free survival (HR=1.65 for [1.08-2.53]; p=0.019) (Supplementary Fig. 2, top panel), as previously observed [12]. Similarly, high *SPARC* mRNA level tended to be associated with shorter recurrence-free survival (HR=1.6 [0.91-2.79]; p=0.097) (Supplementary Fig. 2, bottom panel). We then examined SPARC and cath-D expression by immunohistochemistry (IHC) analysis in serial sections of a TNBC Tissue Micro-Array (TMA) (Fig. 3A). Cath-D was expressed mainly in cancer cells, and to a lesser extent in macrophages, fibroblasts and adipocytes in the tumor stroma (Fig. 3A, left panel). Conversely, SPARC was expressed mainly in fibroblasts, macrophages and endothelial cells, whereas its expression level in cancer cells was variable (Fig. 3A, middle and right panels). Next, we analyzed SPARC and cath-D expression and secretion in five TNBC cell lines and in HMFs (Fig. 3B). Cath-D was expressed by TNBC cell lines and HMFs (Fig. 3B, left panel), but was secreted only by TNBC cells (Fig. 3B, right panel). Conversely, SPARC was expressed and secreted by HMFs, but only by two of the five TNBC cell lines (SUM159 and HS578T) (Fig. 3B). Finally, we investigated SPARC and cath-D co-localization in a TNBC patient-derived xenograft (PDX B1995) [41] in which cath-D expression was previously demonstrated [12]. Co-labelling with polyclonal anti-SPARC and monoclonal anti-cath-D antibodies (Fig. 3C) showed that SPARC (in red; panel a) partially co-localized with cath-D (in green; panel b) in the PDX B1995 microenvironment (merge; panel c). Together with previously published data on SPARC [27, 31–35, 42] and cath-D [7-9, 11, 13, 15-17, 19-21, 43] in BC, our results strongly suggest that it is important to investigate the relationship between SPARC and cath-D that are both co-secreted in the TNBC microenvironment.

**Figure 3.**
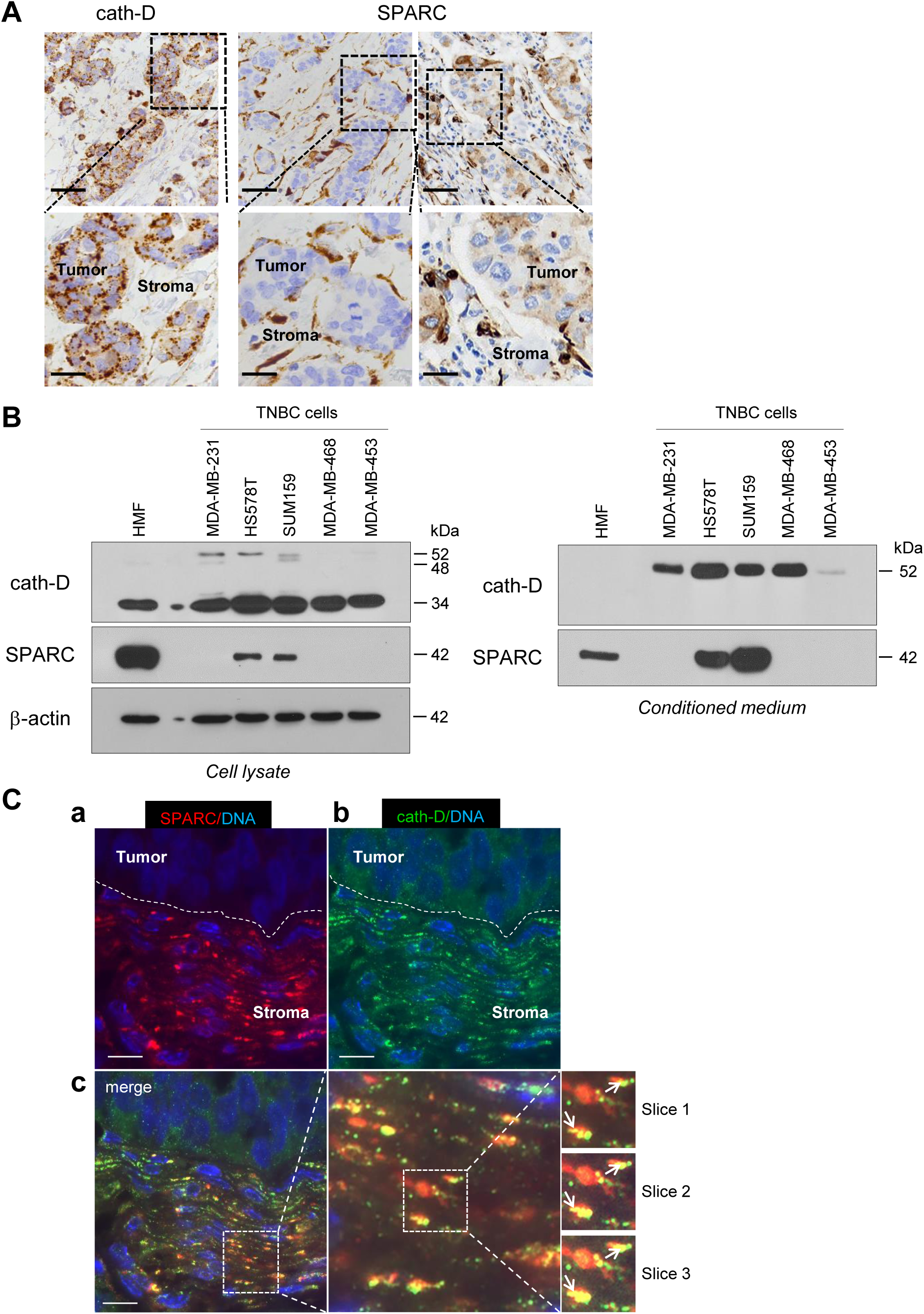
Expression and co-localization of SPARC and cath-D in TNBC. **(A) SPARC and cath-D in a TNBC TMA.** Representative images of SPARC and cath-D expression analyzed in a TNBC TMA (n=147 samples) using anti-SPARC (clone AON-5031) and anti-cath-D (clone C-5) monoclonal antibodies. Scale bars, 50 µm (top panels) and 20 µm (bottom panels; higher magnifications of the boxed regions). **(B) SPARC and cath-D expression and secretion in TNBC cell lines and breast fibroblasts.** Whole cell extracts (10 µg proteins) (left panel) and 24-hour conditioned media in the absence of FCS (40 µl) (right panel) were separated by 13.5% SDS-PAGE and analyzed by immunoblotting with anti-cath-D antibodies for cellular (clone 49, #610801) and secreted cath-D (H-75) detection, respectively, and anti-SPARC (clone AON-5031) antibody. β-actin, loading control. **(C) Co-localization of SPARC and cath-D in TNBC PDX.** PDX 1995 tumor sections were co-incubated with an anti SPARC polyclonal antibody (15274-1-AP) (red; panel a) and an anti-cath-D monoclonal antibody (C-5) (green; panel b). Nuclei were stained with 0.5 µg/ml Hoechst 33342 (blue). Panel c (left): SPARC, cath-D and Hoechst 33342 merge. Panel c (middle and right): higher magnification of the boxed areas (right panels: Z projections of 3 x 0.25 µm slices). Arrows indicate SPARC and cath-D co-localization. Scale bar, 10 µm.

### Cath-D secreted by TNBC cancer cells cleaves fibroblast- and cancer cell-derived SPARC in its extracellular Ca^2+^ binding domain at acidic pH

As the tumor extracellular environment is acidic [44], we then asked whether cath-D can degrade SPARC in the TNBC extracellular medium at low pH. First, we monitored SPARC proteolysis at pH 5.5 in conditioned medium from cath-D-secreting TNBC MDA-MB-231 cells co-cultured with SPARC-secreting HMFs for 24h. SPARC was cleaved in a time-dependent manner in the conditioned medium (Fig. 4A). By western blot analysis, we detected mainly the 34-kDa and 27-kDa SPARC fragments, and to a lesser extent, the 16-kDa fragment (Fig. 4A). Pepstatin A inhibited SPARC cleavage, confirming the involvement of secreted aspartic protease proteolytic activity (Fig. 4A). Moreover, TAILS analysis of the secretome in conditioned medium of co-cultured MDA-MB-231/HMF cells at pH 5.5 showed the presence of the five main SPARC fragments (34-, 27-, 16-, 9-, and 6-kDa) only in the absence of pepstatin A (Table 1, Supplementary Table 1). We then assessed SPARC proteolysis at different pH (6.8 to 5.5), and found that in MDA-MB-231/HMF conditioned medium, SPARC was significantly degraded up to pH 6.2 (Fig. 4B), similarly to the results obtained with recombinant proteins (Fig. 2B). In addition, we observed SPARC limited proteolysis at pH 5.5 also in conditioned medium of HS578T (Fig. 4C) and SUM159 TNBC cells (Fig. 4D) that secrete both proteins. Conversely, we did not observe SPARC cleavage at pH 5.5 in conditioned medium from HMFs co-cultured with MDA-MB-231 cells in which *CTSD* was silenced by RNA interference, indicating that cath-D was responsible for SPARC proteolysis in acidic conditions (Fig. 4E). We confirmed cath-D direct involvement in SPARC proteolysis also by using a mammary cancer cell line derived from tamoxifen-inducible Cre^ERT2,^ *Ctsd^fl/fl^* mice [45] crossed with the transgenic MMTV-PyMT mouse model of metastatic BC [46] (Fig. 4F). In the absence of hydroxytamoxifen (OH-Tam), both cath-D and SPARC were secreted by these cells, whereas cath-D expression and secretion were abrogated by incubation with OH-Tam (Supplementary Fig. 3). SPARC was cleaved in the conditioned medium from this mouse mammary cancer cell line at pH 5.5 only in the absence of OH-Tam (Fig. 4F). These findings demonstrate that cath-D secreted by TNBC and mouse mammary tumor cells cleaves SPARC in its extracellular Ca^2+^ binding domain at the acidic pH found in the tumor microenvironment.

**Figure 4.**
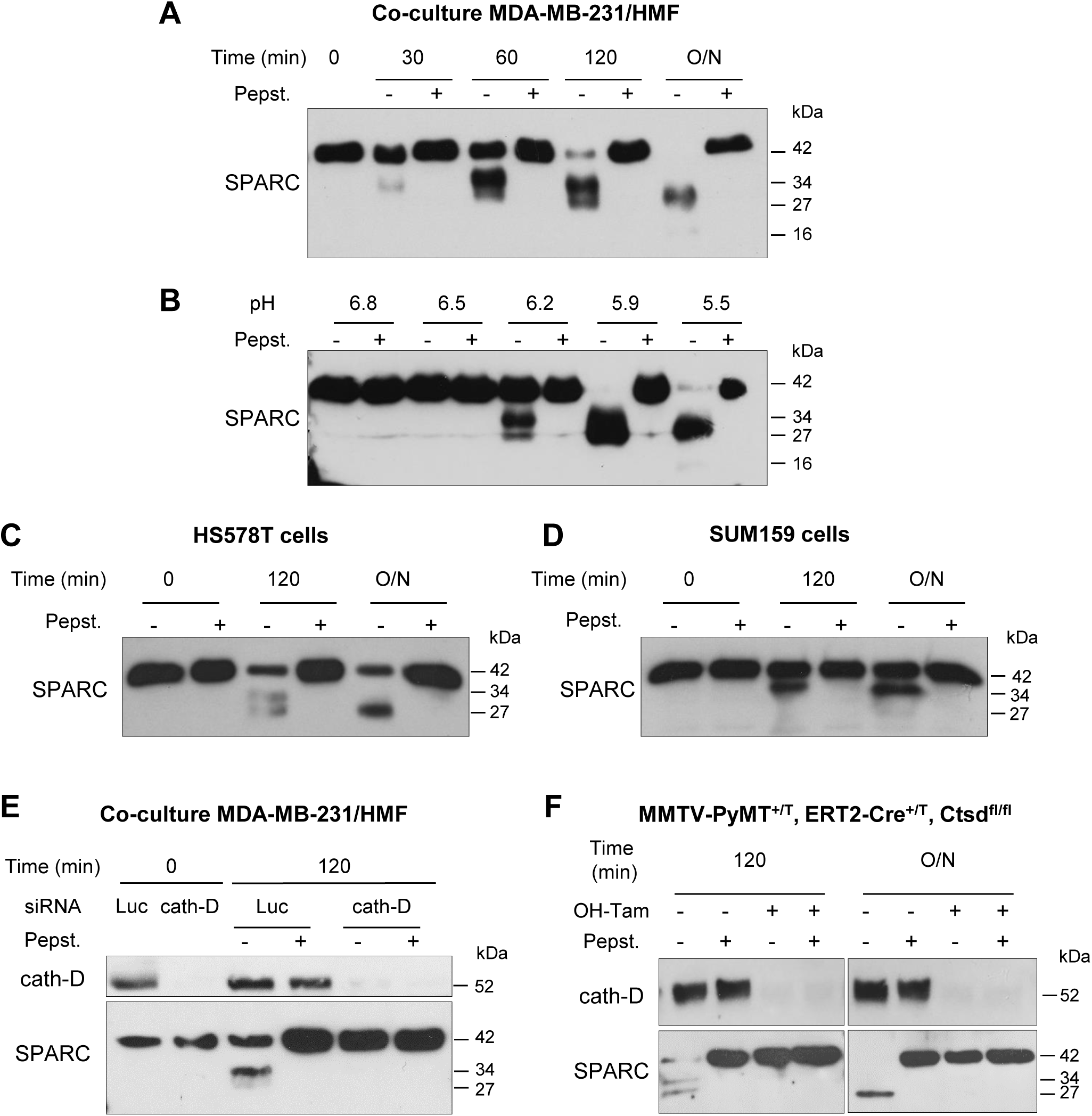
Limited proteolysis of fibroblast- and cancer cell-derived SPARC at acidic pH by cath-D secreted by TNBC and mouse breast cancer cells. **(A) Time-course of SPARC degradation in MDA-MB-231/HMF conditioned medium.** MDA-MB-231 TNBC cells and HMFs were co-cultured in serum-free DMEM without sodium bicarbonate and phenol red and buffered with 50 mM HEPES [pH 7.5] at 37°C for 24h. The 24h conditioned medium from co-cultured MDA-MB-231/HMF was incubated at 37°C in cleavage buffer with or without pepstatin A (Pepst.) at pH 5.5 for the indicated times. SPARC cleavage in conditioned medium was analyzed by 13.5% SDS-PAGE and immunoblotting with an anti-SPARC antibody (15274-1-AP). O/N, overnight. **(B) Influence of the milieu acidity on SPARC degradation in MDA-MB-231/HMF conditioned medium.** MDA-MB-231 TNBC cells and HMFs were co-cultured as in (**A**). The 24h conditioned medium was incubated at 37°C in cleavage buffer with or without pepstatin A at the indicated pH overnight. SPARC cleavage was analyzed as described in (**A**). **(C and D) Time-course of SPARC cleavage in in TNBC cell conditioned medium.** HS578T TNBC cells **(C)** and SUM159 TNBC cells **(D)** were cultured in serum-free DMEM without sodium bicarbonate and phenol red and buffered with 50 mM HEPES [pH 7.5] at 37°C for 24h.The 24h conditioned medium was incubated at 37°C in cleavage buffer with or without pepstatin A at pH 5.5 for the indicated times. SPARC cleavage was analyzed as described in (**A**). **(E) SPARC cleavage by cath-D secreted by MDA-MB-231 cells.** MDA-MB-231 cells were transfected with Luc or cath-D siRNAs. At 48h post-transfection, siRNA-transfected MDA-MB-231 cells were co-cultured with HMFs as described in (**A**). Then, the 24h conditioned media from co-cultured siRNA-transfected MDA-MB-231/HMF were incubated at 37°C in cleavage buffer with or without pepstatin A at pH 5.5 for 120 min. Cath-D secretion by siRNA-transfected MDA-MB-231 cells was analyzed with an anti-cath-D antibody (H-75). SPARC cleavage was analyzed as described in (**A**). **(F) SPARC cleavage by cath-D secreted by inducible *Ctsd* knock-out MMTV-PyMT mammary tumor cells.** Inducible *Ctsd* knock-out MMTV-PyMT breast cancer cells were incubated or not with 4-hydroxytamoxifen (OH-Tam; 3 µM) for 4 days to induce *Ctsd* knock-out. Then, cells were cultured in FCS-free DMEM without sodium bicarbonate and phenol red and buffered with 50 mM HEPES [pH 7.5] at 37°C for 24h. The 24h-conditioned medium was incubated at 37°C in cleavage buffer with or without pepstatin A at pH 5.5 for 120 min or O/N. Cath-D secretion was analyzed with an anti-cath-D antibody (AF1029). SPARC cleavage was analyzed as described in (**A**).

### SPARC is cleaved *in vivo* in TNBC and mouse mammary tumors

To demonstrate that SPARC is cleaved by cath-D also *in vivo*, we first analyzed the level of FL SPARC and cleaved fragments in whole cytosols of mammary tumors from MMTV-PyMT *Ctsd^-/-^* knock-out mice (Fig. 5A). As expected, cath-D was expressed in the cytosol of mammary tumors from MMTV-PyMT *Ctsd^+/+^*, but not from MMTV-PyMT, *Ctsd^-/-^* mice (Fig. 5A, left panels). Moreover, in two of the three tumors from MMTV-PyMT, *Ctsd^-/-^* mice, SPARC expression level was much higher than in the three tumors from MMTV-PyMT, *Ctsd^+/+^* mice (Fig. 5A, left panel). Unexpectedly, we could not detect any SPARC cleavage fragment in this transgenic mouse model, certainly due to further SPARC proteolysis *in vivo* by other proteinases. Nevertheless, SPARC reduction occurred through post-translational mechanisms because *Sparc* mRNA level was not significantly different in the corresponding MMTV-PyMT, *Ctsd^+/+^* and MMTV-PyMT, *Ctsd^-/-^* tumors (Fig. 5A, right panel). We then evaluated the presence of FL SPARC and its cleaved fragments in the whole cytosols from two TNBC PDXs that express cath-D at high and low level, respectively (Fig. 5B, top panel). We detected FL SPARC and its 34-kDa cleaved fragment in PDX B3977 (high cath-D expression), but only FL SPARC in PDX B1995 (low cath-D expression) (Fig. 5B, bottom panel). Finally, analysis of whole cytosols from two clinical TNBC samples with different cath-D expression levels (Fig. 5C, top panel) showed that FL SPARC level was lower in cytosol C1 (TNBC with high cath-D expression) than in cytosol C2 (TNBC with low cath-D expression) (Fig. 5C, bottom panel). Moreover, we detected the 27-kDa cleaved SPARC fragment only in cytosol C1 (Fig. 5C, bottom panel). Overall, these results strongly suggest that SPARC cleavage in its extracellular Ca^2+^ binding domain may occur *in vivo* in cath-D-expressing mammary cancers, although other proteinases may also be involved.

**Figure 5.**
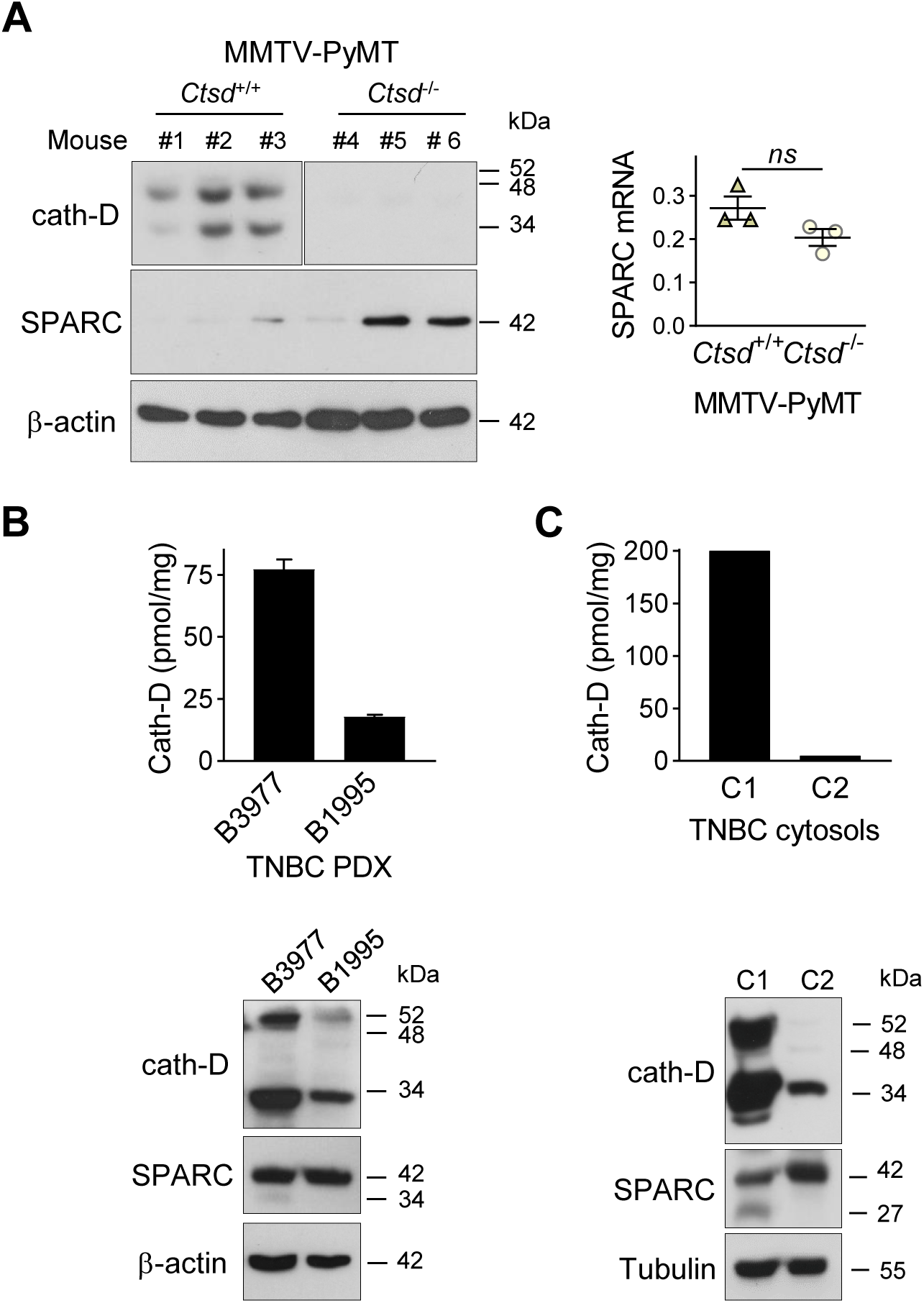
Detection of FL SPARC and its cleaved fragments in mammary tumors. **(A) SPARC expression in mammary tumors from MMTV-PyMT *Ctsd* knock-out mice**. Left panel, whole cytosols (40 µg) of mammary tumors from MMTV-PyMT*^Ctsd+/+^* (N°1-3) and MMTV-PyMT*^Ctsd-/-^* (*Ctsd* knock-down in mammary glands) (N°4-6) mice were analyzed by 13.5% SDS-PAGE and immunoblotting with anti-mouse cath-D (clone 49, #610801) and anti-SPARC (AON-5031) antibodies. β-actin, loading control. Right panel, total RNA was extracted from mammary tumors from MMTV-PyMT*^Ctsd+/+^* (N°1-3) and MMTV-PyMT*^Ctsd-/-^* (N°4-6) mice, and *Sparc* expression was analyzed by RT-qPCR. *P*=0.1 (Student’s *t*-test). **(B and C) SPARC expression in TNBC PDXs and TNBC biopsies**. Top panels, cath-D expression was determined in whole cytosols from two TNBC PDXs **(B)** and two TNBC biopsies **(C)** by sandwich ELISA with the immobilized anti-human cath-D D7E3 antibody and the anti-human cath-D M1G8 antibody coupled to HRP. Bottom panels, whole cytosols (40 µg) from these PDXs **(B)** and TNBC biopsies **(C)** were analyzed by 13.5% SDS-PAGE and immunoblotting with anti-cath-D (H-75) and anti-SPARC (15274-1-AP) antibodies. β-actin **(B)** and tubulin **(C)**, loading controls.

### Cath-D-induced SPARC fragments inhibit TNBC cell adhesion and spreading on fibronectin, and promote their motility, endothelial transmigration and invasion

Previous studies reported that FL SPARC and particularly its C-terminal extracellular Ca^2+^ binding domain can modulate adhesion, spreading, motility, endothelial transmigration, and invasion of cancer and stromal cells [25, 26, 33, 47–49]. Therefore, we compared the effect of the cath-D-induced SPARC fragments (mixture of 34+27+16+9+ 6-kDa fragments) (Supplementary Fig. 4) and of FL recombinant SPARC (42-kDa) in MDA-MB-231 cells. Soluble FL SPARC significantly inhibited MDA-MB-231 cell adhesion on fibronectin in a dose-dependent manner (Supplementary Fig. 5). After incubation with FL SPARC (final concentration of 10 µg/ml, 240 nM), as previously described [26, 47], MDA-MB-231 cell adhesion on fibronectin was reduced by 1.3-fold compared with control (CTRL; untreated) (Fig. 6A; *P<0.001).* Moreover, FL SPARC inhibition of cell adhesion was similar in Luc- and cath-D-silenced MDA-MB-231 cells, indicating an autonomous effect of SPARC on cell adhesion (Supplementary Fig. 6). Incubation of MDA-MB-231 cells with cath-D-induced SPARC fragments (cleaved SPARC) also significantly decreased cell adhesion by 1.7-fold compared with control (CTRL) (Fig. 6A; *P<0.001)* and by 1.3-fold compared with FL SPARC (Fig. 6A; *P<0.001).* We also monitored the effects of FL and cleaved SPARC on MDA-MB-231 cell spreading on fibronectin by staining F-actin filaments with phalloidin (Supplementary Fig. 7A). Both FL SPARC and SPARC cleaved fragments led to a decrease of the cell surface contact area on fibronectin through F-actin peripheral rearrangement. Specifically, bundling of actin stress fibers was disrupted and actin microfilaments were redistributed in a peripheral web (Supplementary Fig. 7A). This suggests a transition to an intermediate state of adhesiveness, previously described for FL SPARC [50], that may favor cell migration and invasion [51]. Incubation with FL and cleaved SPARC decreased the percentage of spread cells by 2.1-fold and 3.8-fold, respectively, compared with control (Supplementary Fig. 7B; *P<0.001*). This inhibition was significantly higher (1.8-fold) with cleaved SPARC than FL SPARC (Supplementary Fig. 7B; *P<0.05*). Then, cell motility analysis in Boyden chambers showed quite high basal motility of MDA-MB-231 cells, as expected for mesenchymal cells (e.g. 49% of cells passed through the fibronectin-coated filters) (Fig. 6B). Incubation with FL and cleaved SPARC increased MDA-MB-231 cell motility by 1.5-fold and 1.9-fold, respectively, compared with control (Fig. 6B; *P<0.01* and *P<0.001).* Moreover, the effect of cleaved SPARC on cell motility was 1.3-fold higher than that of FL SPARC (Fig. 6B; *P<0.05*). In the endothelial transmigration assay, FL and cleaved SPARC fragments stimulated MDA-MB-231 migration through primary human umbilical vein endothelial cells (HUVECs) by 1.4-fold and 1.7-fold, respectively, compared with control (Fig. 6C; *P<0.01* and *P<0.001*). The effect of cleaved SPARC was 1.2-fold higher than that of FL SPARC (Fig. 6C; *P<0.05*). Finally, both FL and cleaved SPARC fragments increased MDA-MB-231 cell invasion through Matrigel-coated filters in Boyden chambers by 2-fold and 3-fold, respectively, compared with control (Fig. 6D; *P<0.001).* The effect of cleaved SPARC was 1.5-fold higher than that of FL SPARC (Fig. 6D; *P<0.001).* Altogether, these results indicate that FL SPARC inhibits MDA-MB-231 cell adhesion and spreading, and promotes MDA-MB-231 cell motility, endothelial transmigration, and invasion. These effects were increased by incubation with cath-D-induced SPARC fragments, suggesting that in the TNBC microenvironment, cath-D amplifies SPARC pro-tumor activity through proteolysis of its extracellular Ca^2+^ binding domain.

**Figure 6.**
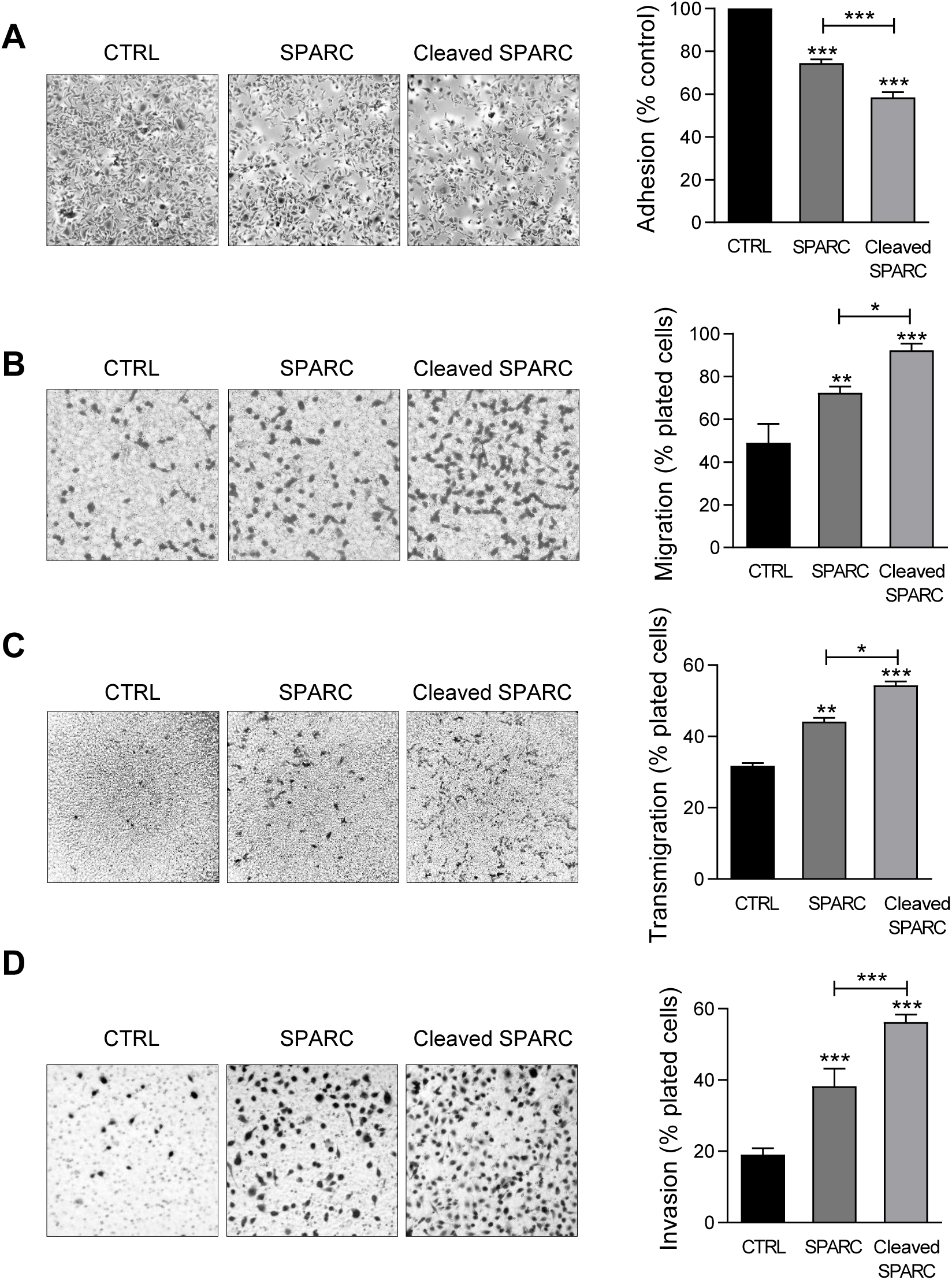
Effects of FL SPARC and cath-D-induced cleaved SPARC fragments on adhesion, migration, transmigration and invasion of TNBC cells. **(A) Cell adhesion.** MDA-MB-231 cells were let to adhere for 30 min on a fibronectin matrix in the absence or presence of recombinant FL SPARC (SPARC), or recombinant cath-D-induced cleaved SPARC fragments (cleaved SPARC) at the final concentration of 240 nM. Left panels, representative images of adherent cells. Right panel, adherent cells were stained with crystal violet, and adhesion was quantified at 570 nm. CTRL, PBS in cleavage buffer. Data are the mean ± SD (n=3); ***, p<0.001, ANOVA and Bonferroni’s post hoc test. Similar results were obtained in four independent experiments. **(B) Cell migration.** MDA-MB-231 cells were let to migrate for 16h on a fibronectin matrix in the absence or presence of recombinant FL SPARC, or cleaved SPARC at a final concentration of 240 nM. Left panels, representative images of migrating cells. Right panel, migrating cells were quantified by MTT staining and absorbance was read at 570 nm. CTRL, PBS in cleavage buffer. Data are the mean ± SD (n=3); *, p<0.05; **, p<0.01; ***, p<0.001, ANOVA and Bonferroni’s post hoc test. Similar results were obtained in three independent experiments. **(C) Endothelial transmigration.** MDA-MB-231 cells were let to transmigrate for 16h through a HUVEC monolayer in the absence or presence of recombinant FL SPARC, or cleaved SPARC at a final concentration of 240 nM. Left panels, representative images of transmigrating cells. Right panel, transmigrating cells were stained with MTT and quantified at 570 nm. CTRL, PBS in cleavage buffer. Data are the mean ± SD (n=3); *, p<0.05, **, p<0.01, ***, p<0.001, ANOVA and Bonferroni’s post hoc test. Similar results were obtained in two independent experiments. **(D) Cell invasion.** MDA-MB-231 cells were let to invade for 16h on a Matrigel matrix in the absence or presence of recombinant FL SPARC, or cleaved SPARC at a final concentration of 240 nM. Left panels, representative images of invading cells. Right panel, invading cells were stained with MTT and quantified at 570 nm. CTRL, PBS in cleavage buffer. Data are the mean ± SD (n=3); ***, p<0.001, ANOVA and Bonferroni’s post hoc test. Similar results were obtained in three independent experiments.

### The 9-kDa C-terminal SPARC fragment inhibits TNBC cell adhesion and spreading on fibronectin, and promotes their motility, endothelial transmigration, and invasion

To identify the SPARC domain(s) involved in these functions, we produced FL SPARC and its various cleaved fragments in mammalian cells and purified them, as previously described [52, 53] (Fig. 7A, Supplementary Table 2). We first determined which SPARC fragment(s) were involved in the reduction of cell adhesion by incubating MDA-MB-231 cells with equimolar amounts of FL protein and each fragment (Fig. 7B). As before (Fig. 6A), purified FL SPARC (42-kDa) reduced MDA-MB-231 cell adhesion by 1.4-fold compared with control (Fig. 7B; *P<0.001*). However, among the C-terminal SPARC fragments, only the 9-kDa fragment (amino acids 235-303) significantly decreased MDA-MB-231 cell adhesion by 2-fold compared with control (Fig. 7B; *P<0.001*), and by 1.4-fold compared with FL SPARC (Fig. 7B; *P<0.001*). The 9-kDa C-terminal SPARC fragment (amino acids 235-303) contains the two Ca^2+^ binding sequences of the two EF-hand domains (Supplementary Fig. 8) that are involved in focal adhesion disassembly, and are crucial for SPARC-mediated inhibition of adhesion [47, 54]. The 16-kDa C-terminal SPARC fragment (amino acids 179-303) reduced cell adhesion by 1.2-fold (not significant) (Fig. 7B and Supplementary Fig. 8), and the 6-kDa SPARC fragment (amino acids 258-303) had no effect (Fig. 7B and Supplementary Fig. 8). Therefore, among the five cath-D-induced SPARC fragments (Fig. 2E), only the C-terminal 9-kDa fragment could inhibit cell adhesion and more potently than FL SPARC.

**Figure 7.**
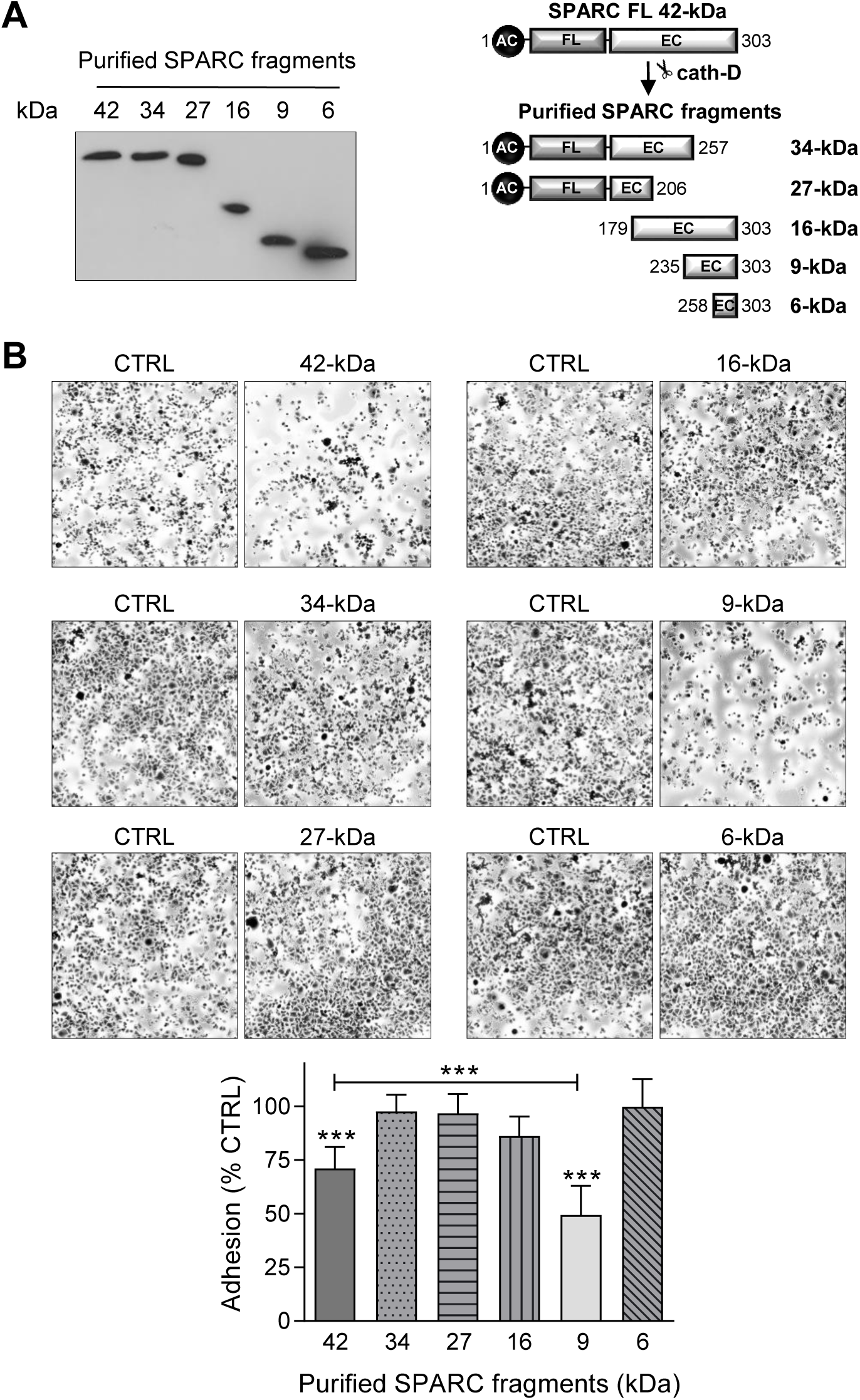
Effect of FL SPARC and cath-D-induced cleaved SPARC fragments on TNBC cell adhesion. **(A) Production of Myc/His-tagged FL SPARC, and Myc/His-tagged 34-, 27-, 16-, 9-, and 6-kDa SPARC fragments.** Left panel, equimolar concentrations (240 nM each) of purified Myc/His-tagged FL SPARC and SPARC fragments were analyzed by SDS-PAGE (17%) and immunoblotting with an anti-Myc antibody (clone 9B11). Right panel, schematic representation of the purified Myc/His-tagged SPARC fragments. AC, acidic domain; FL, follistatin-like domain; EC, Ca^2+^-extracellular binding domain. **(B) Cell adhesion.** MDA-MB-231 cells were let to adhere for 30 min on a fibronectin matrix in the presence of purified Myc/His-tagged FL SPARC, or individual Myc/His-tagged SPARC fragments (34-, 27-, 16-, 9-, and 6-kDa) at an equimolar final concentration (240 nM each). Upper panels, representative images of adherent cells stained with crystal violet after incubation with the indicated SPARC variants. Lower panel, cell adhesion was quantified as described in Fig. 6 A and expressed as percentage relative to the value in control (SPARC-immunodepleted control for each SPARC fragment). Data are the mean ± SD of three independent experiments; ***, p<0.001, ANOVA and Bonferroni’s post hoc test.

Based on these results, we compared the effects on MDA-MB-231 cell adhesion, spreading, motility, endothelial transmigration and invasion of the 9-kDa C-terminal SPARC fragment, FL SPARC, and cath-D-induced SPARC fragments (mixture of 34+27+16+9+ 6-kDa fragments) (Fig. 8, Supplementary Figs. 9, and 10). Incubation with the 9-kDa C-terminal SPARC fragment significantly decreased cell adhesion (Fig. 8A, *P<0.001*) and spreading (Supplementary Fig. 10, *P<0.001*) by 2.1-fold and 8.4-fold, respectively, compared with control, and significantly increased cell motility by 1.6-fold (Fig.8B; *P<0.001)*, endothelial transmigration by 2.1-fold (Fig. 8C; *P<0.001),* and cell invasion by 1.7-fold (Fig. 8D; *P<0.001)* compared with control. Moreover, the 9-kDa SPARC fragment seemed to induce a transition to an intermediate adhesive state highlighted by the loss of actin-containing stress fibers (Supplementary Fig. 10A). Compared with FL SPARC, incubation with the 9-kDa C-terminal SPARC fragment significantly decreased cell adhesion (Fig. 8A, *P<0.001*) and spreading (Supplementary Fig. 10, *P<0.001*) by 1.7-fold and 4.2-fold, respectively, and significantly increased cell motility by 1.2-fold (Fig.8B; *P<0.001)*, endothelial transmigration by 1.3-fold (Fig. 8C; *P<0.05),* and cell invasion by 1.4-fold (Fig. 8D; P<0.01). Conversely, we did not observe any significant difference between the 9-kDa C-terminal SPARC and the cath-D-induced SPARC fragments (Fig. 8 and Supplementary Fig. 10). These findings demonstrate that the 9-kDa C-terminal SPARC released by cath-D exhibits pro-tumor activity in the TNBC microenvironment.

**Figure 8.**
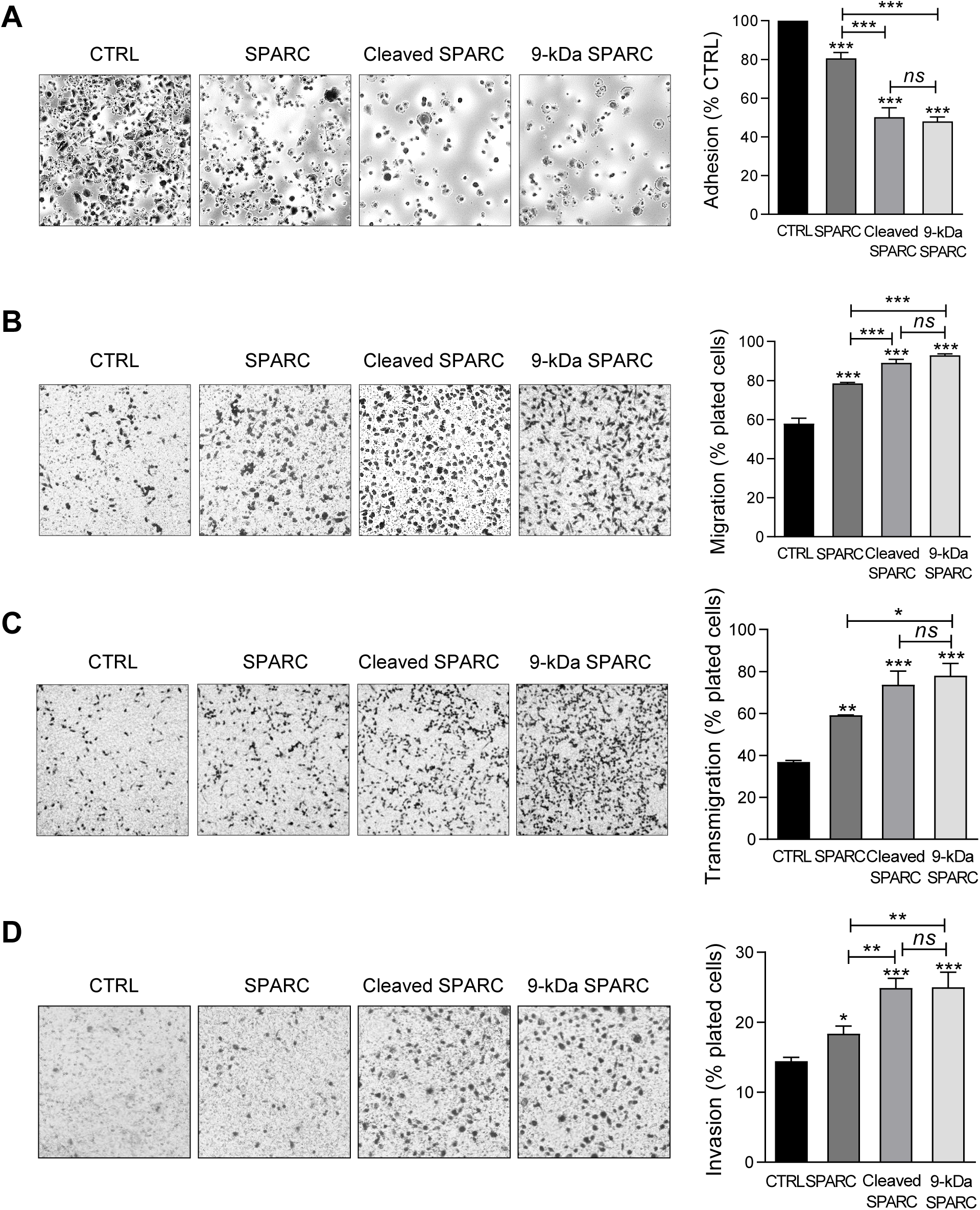
Effects of the 9-kDa C-terminal SPARC fragment on TNBC cell adhesion, migration, transmigration and invasion. **(A) Cell adhesion.** MDA-MB-231 cells were let to adhere for 30 min on a fibronectin matrix in the presence of recombinant FL SPARC, recombinant cleaved SPARC fragments (cleaved SPARC), or purified 9-kDa C-terminal SPARC fragment at a final concentration of 240 nM. Left panels, representative images of adherent cells stained with crystal violet. Right panel, adhesion was quantified as described in Fig. 6 A. Data are the mean ± SD (n=3); ns, not significant; ***, p<0.001, ANOVA and Bonferroni’s post hoc test. CTRL, PBS in cleavage buffer and SPARC-immunodepleted supernatant from the 9-kDa SPARC fragment purification. Similar results were obtained in three independent experiments. **(B) Cell migration.** MDA-MB-231 cells were let to migrate for 16h on a fibronectin matrix in the absence or presence of FL SPARC, cleaved SPARC fragments, or the 9-kDa C-terminal SPARC fragment at a final concentration of 240 nM. Left panels, representative images of migrating cells stained with crystal violet. Right panel, migration was quantified as described in Fig. 6 B. Data are the mean ± SD (n=3); ***, p<0.001, ANOVA and Bonferroni’s post hoc test. CTRL, PBS in cleavage buffer and SPARC immunodepleted supernatant from the 9-kDa SPARC fragment purification. Similar results were obtained in two independent experiments. **(C) Endothelial transmigration.** MDA-MB-231 cells were let to transmigrate for 16h through a HUVEC monolayer in the absence or presence of FL SPARC, cleaved SPARC fragments, or the 9-kDa C-terminal SPARC fragment at a final concentration of 240 nM. Left panels, representative images of transmigrating cells. Right panel, transmigrating cells were stained with MTT and quantified by absorbance at 570 nm. Data are the mean ± SD (n=3); *, p<0.05, **, p<0.01, ***, p<0.001, ANOVA and Bonferroni’s post hoc test. CTRL, PBS in cleavage buffer and SPARC-immunodepleted supernatant from the 9-kDa SPARC fragment purification. Similar results were obtained in two independent experiments. **(D) Cell invasion.** MDA-MB-231 cells were let to invade for 16h on a Matrigel matrix in the absence or presence of FL SPARC, cleaved SPARC fragments, or the 9-kDa C-terminal SPARC fragment at a final concentration of 240 nM. Left panels, representative images of invading cells stained with crystal violet. Right panel, invading cells were quantified by absorbance at 570 nm. Data are the mean ± SD (n=3); *, p<0.05, **, p<0.01, ***, p<0.001, ANOVA and Bonferroni’s post hoc test. CTRL, PBS in cleavage buffer and SPARC immunodepleted supernatant from the 9-kDa SPARC fragment purification. Similar results were obtained in two independent experiments.

## DISCUSSION

This study shows that cath-D secreted by TNBC cells triggers fibroblast- and cancer cell-derived SPARC cleavage at the acidic pH of the tumor microenvironment, leading to the production of the bioactive 9-kDa C-terminal SPARC fragment that inhibits cancer cell adhesion and spreading on fibronectin, and stimulates their migration, endothelial transmigration and invasion (Fig. 9). The TAILS analysis of the secretome of conditioned medium from TNBC cells co-cultured with HMFs revealed that five main SPARC fragments (34-, 27-, 16-, 9-, and 6-kDa) are released in the extracellular environment at acidic pH. Our previous TAILS-based study showed that cystatin C is a substrate of extracellular cath-D and it is completely degraded by multiple cleavage, highlighting the complexity of the proteolytic cascades that operate in the tumor microenvironment [55]. Here, we demonstrate that cath-D triggers also the limited proteolysis of the matricellular protein SPARC in an acidic environment to favor TNBC invasion.

**Figure 9.**
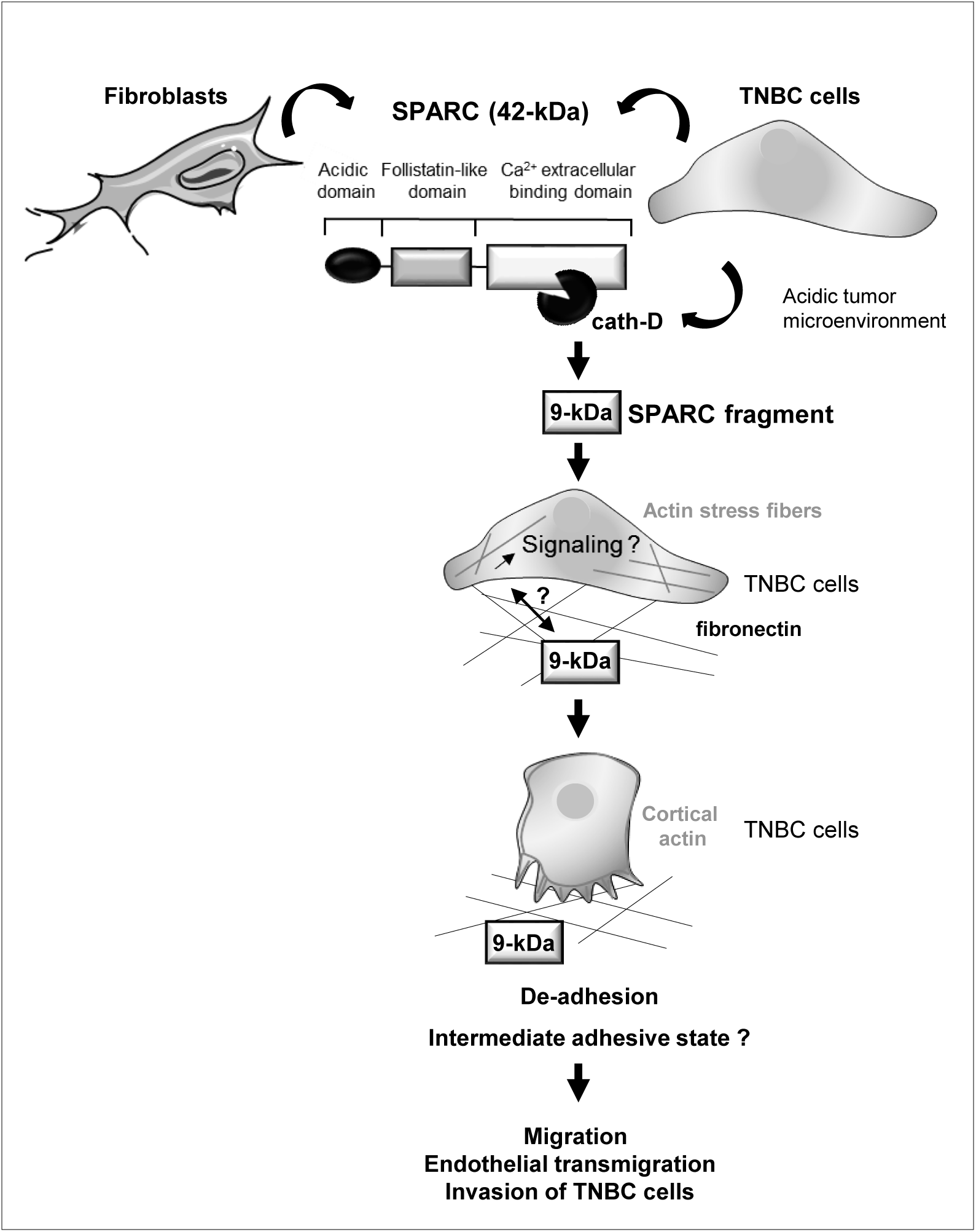
Model of the pro-tumor effect on TNBC cells of the 9-kDa C-terminal SPARC released by cath-D cleavage. TNBC-secreted cath-D triggers limited proteolysis of SPARC at the acidic pH of the tumor microenvironment. Among the SPARC fragments cleaved by cath-D, the 9-kDa C-terminal SPARC fragment inhibits TNBC cell adhesion and spreading. This might lead to an intermediate adhesive state, and stimulate TNBC cell migration, endothelial transmigration and invasion.

Our recent study indicated that extracellular cath-D is a therapeutic target for immunotherapy and a TNBC biomarker [12]. Moreover, Huang *et al* found that cath-D is overexpressed in 71.5% of the 504 TNBC samples analyzed, and proposed a prognostic model for TNBC outcome based on node status, cath-D expression, and KI67 index [10]. Furthermore, co-expression of cath-D and androgen receptor defines a TNBC subgroup with poorer overall survival [11]. SPARC protein and mRNA are overexpressed in TNBC [34, 36, 56], and this has been associated with poor prognosis [33, 34, 36]. Here, we showed that high *CTSD* and *SPARC* mRNA expression tended to be associated with shorter recurrence-free survival in a cohort of 255 patients with TNBC using an on line survival tool [40]. Moreover, in a TNBC TMA, we found that cath-D was mainly expressed by cancer cells and some stromal cells, as shown previously [12]. Conversely, SPARC was mainly expressed in mesenchymal cells, while its expression level in tumor cells was variable, as previously described [27, 28]. *In cellulo,* cath-D was secreted by TNBC cells and SPARC by human breast fibroblasts and some TNBC cell lines, as previously described [11, 38, 42]. Importantly, cath-D and SPARC co-localized in the microenvironment of TNBC PDX. Overall, these data prompted us to study the interplay between cath-D and SPARC in the TNBC microenvironment.

We then demonstrated that cath-D cleaves SPARC *in vitro* in an acidic environment exclusively in its extracellular Ca^2+^ binding domain, specifically releasing five main SPARC fragments (34-, 27-, 16-, 9-, and 6-kDa). The main peptide bonds cleaved by cath-D at low pH are Phe-Phe, Leu-Tyr, Tyr-Leu, and Phe-Tyr [57] that correspond relatively well to the cleavage sites identified in this study. Interestingly, the presence of additional cleavage sites (e.g. Leu-Val, Leu-Asp/Glu or Gln-Phe, Gly-Tyr and Ala-Pro), which confirms cath-D preference for cleavage sites with at least one hydrophobic residue in P1 or P1’. SPARC biological activity can be modulated by limited proteolysis, leading to the unmasking of distinct or amplified biological functions compared with those of the FL protein [23, 49]. For instance, matrix metalloproteinases (MMP-1, -2, -3, -9 and -13) cleave SPARC *in vitro* in its N-terminal acid domain and in its extracellular Ca^2+^ binding domain, releasing fragments with higher affinity for collagens that modulate cell-cell and cell-matrix extracellular interactions in the tumor microenvironment [58]. In addition, MMP-3-mediated SPARC cleavage *in vitro* produces fragments that affect angiogenesis [59]. More recently, cleavage of SPARC extracellular Ca^2+^ binding domain by MMP-8 and MMP-13 has been detected in the serum of patients with lung cancer, indicating their presence also *in vivo* [60]. Similarly, the cysteine cathepsin K (cath-K) also cleaves SPARC *in vitro* and *in vivo* in its N-terminal acid domain, and in its extracellular Ca^2+^ binding domain in prostate cancer bone metastases, releasing a 10-kDa C-terminal fragment with unknown biological activity [61]. The 9-kDa SPARC fragment generated by cath-D in our study is within the 10-kDa SPARC fragment generated by cath-K [61].

We then demonstrated that at acidic pH, cath-D present in conditioned medium from cath-D-secreting MDA-MB-231 TNBC cells co-cultured with SPARC-secreting HMFs (or from HS578T and SUM159 TNBC cells that secrete both factors) induces limited proteolysis of SPARC, leading to the production of SPARC fragments with the same molecular weight (34-, 27-, and 16-kDa), as detected with the recombinant proteins. By western blot analysis, we could not detect the 9-kDa and 6-kDa SPARC fragments, whereas we identified all five SPARC main fragments (34-, 27-, 16-, 9- and 6-kDa) in the TAILS analysis of the secretomes of MDA-MB-231/HMF co-cultures. Moreover, in breast cancers of transgenic mice, in TNBC PDXs, and in two clinical TNBC samples that express high or low cath-D levels, we found that cath-D expression was inversely correlated with SPARC levels, and detected the SPARC fragments of 34-kDa and 27-kDa only in samples that strongly expressed cath-D. Altogether, our data strongly suggest that cath-D cleaves SPARC in its extracellular Ca^2+^ binding domain *in vitro*, in the secretome of TNBC cell lines, and *in vivo* in TNBC. Importantly, SPARC peptides corresponding to the 27-kDa (207-218), 16-kDa (179-191), 9-kDa (236-256), and 6-kDa (258-268) fragments have been found in BC samples by using an on line proteomic database (https://cptac-data-portal.georgetown.edu/) [62]. To date, only one study on cath-K in prostate cancer [61] validated *in vitro, in cellulo* and *in vivo* SPARC cleavage events by proteases in cancer. In addition, to our knowledge, no study has established yet a direct link between a SPARC fragment generated by a protease activity and its biological function in TNBC.

SPARC plays multiple contextual functions depending on the cancer type and stage, and its precise role(s) in TNBC remains to be studied. FL SPARC stimulates migration and invasion of TNBC cells [63], and promotes MMP-2 activation in TNBC cells, thereby contributing to the proteolytic cascades associated with tumor invasion [42]. In addition, FL SPARC stimulates tumor growth and lung colonization after grafting of mouse TNBC 4T1 and LM3 cells in syngeneic mice by promoting cell cycling and expansion of myeloid-derived suppressor cells (MDSCs) [35]. Conversely, FL SPARC transfected in high-grade isogenic BC cells reduces tumor rate, and favors epithelial-to-mesenchymal transition and the formation of a highly immunosuppressive microenvironment composed of immune cells, such as MDSCs [33]. However, the oncogenic roles of SPARC fragments generated by proteolysis in the tumor microenvironment of TNBC were not known. Here, we demonstrated that the 9-kDa C-terminal fragment of SPARC released by cath-D has greater oncogenic activity than FL SPARC, showing a new crosstalk between proteases and matricellular proteins in the TNBC stroma. We found that the 9-kDa C-terminal fragment of SPARC is an important regulator of MDA-MB-231 TNBC cell adhesion, spreading, migration, endothelial transmigration and invasion. Specifically, it inhibited cell adhesion and spreading on fibronectin, and induced redistribution of actin microfilaments at the periphery of TNBC cells associated with cell rounding. A previous work showed that FL SPARC inhibits endothelial cell spreading and induces endothelial cell rounding by affecting the early stages of the counter-adhesive process through the loss of vinculin-containing focal adhesion plaques and the concomitant reorganization of actin-containing stress fibers [47]. This might correspond to an intermediate adhesive state that was previously observed in endothelial cells incubated with FL SPARC [64], and in some cancer types, such as gliomas [65], and that promotes cell motility and invasion [51].

The 9-kDa C-terminal fragment is located in the extracellular Ca^2+^ binding domain of SPARC. The crystal structure of this SPARC domain (Protein Data Bank DOI: 10.2210/pdb2V53/pdb) shows a canonical pair of EF-hand calcium binding sites that are essential for stabilizing Ca^2+^ binding [66]. Both amino- and carboxyl-terminal (EF-hand) domains of SPARC bind to Ca^2+^ that is required for maintenance of its native structure [54]. In the EF-hand motif, two helices (E and F) flank a loop of 12 amino acids in which the Ca^2+^ ion is coordinated in a pentagonal bipyramidal arrangement [67]. Moreover, FL SPARC binding to the extracellular matrix is Ca^2+^-dependent [54]. Interestingly, synthetic small peptides (^272^TCDLDNDKYIALDEWAGCFG^291^) with sequences derived from SPARC C-terminal extracellular Ca^2+^ binding domain (EF hand-2) inhibit adhesion and spreading of endothelial cells and fibroblasts [47, 50, 68]. In our experimental model, it seems unlikely that only the EF hand-2 (aa 262-294) domain is involved in inhibiting MDA-MB-231 cell adhesion to fibronectin. Indeed, the 6-kDa SPARC fragment (amino acids 258-303) that contains only the EF hand-2 domain did not inhibit MDA-MB-231 cell adhesion, unlike the 9-kDa SPARC fragment (amino acids 235-303) that contains the residues coordinating Ca^2+^ in both EF-hands. This suggests that the two Ca^2+^-binding domains are involved in this effect. However, inhibition of cell adhesion by the 16-kDa SPARC fragment (amino acids 179-303) that contains both EF-hand domains was less important compared with the 9-kDa SPARC fragment. This suggests that the additional N-terminal sequences may alter the EF-hand domain conformation, or may interfere with Ca^2+^ binding or with the interaction with fibronectin or a TNBC cell surface receptor. Moreover, the three-dimensional conformation of the N-terminal sequences of the 16-kDa fragment might be different from that of FL SPARC, which significantly inhibited MDA-MB-231 cell adhesion. It remains to be determined whether the 9-kDa C-terminal fragment of SPARC acts directly through a specific receptor, such α5β1 integrin, as described for FL SPARC [25], or by blocking adhesive interactions.

Finally, we demonstrated that FL SPARC and more strongly the 9-kDa C-terminal fragment promoted TNBC cell endothelial transmigration, an essential step for extravasation and metastasis. Similarly, a previous study showed that the C-terminal extracellular Ca^2+^ module of SPARC, a domain implicated in binding to endothelial cells [69] and to vascular cell adhesion molecule 1 (VCAM1) [70], is needed to enhance endothelial transmigration of melanoma cells *via* VCAM1 signaling [26]. These findings suggest a role for the 9-kDa C-terminal SPARC fragment in vascular permeability, extravasation and metastasis formation *in vivo*.

Our current results indicate that cath-D secreted by TNBC cells is part of the proteolytic network in the TNBC acidic microenvironment that generates a bioactive 9-kDa C-terminal fragment of the matricellular protein SPARC with enhanced oncogenic activity. We dissected the molecular mechanisms that link SPARC limited cleavage by cath-D in TNBC microenvironment to the amplified oncogenic activity of a 9-kDa SPARC C-terminal fragment, highlighting a novel paradigm of alteration of TNBC extracellular milieu by proteolysis. Overall, these results indicate that the 9-kDa C-terminal SPARC fragment is an attractive target for cancer therapies in TNBC, and open the way for developing novel targeted therapies against bioactive fragments from matricellular proteins, for restructuring the surrounding microenvironment and reducing tumorigenesis [4].

## MATERIALS AND METHODS

### Antibodies

The rabbit polyclonal anti-SPARC antibody (15274-1-AP) was purchased from Proteintech. The mouse monoclonal anti-human SPARC (clone AON-5031, sc-73472), the rabbit polyclonal anti-human cath-D antibody (H-75, sc-10725), and the mouse monoclonal anti-human cath-D (clone C-5, sc-377124) antibodies were purchased from Santa Cruz Biotechnology. The mouse monoclonal anti-human cath-D antibody (clone 49, #610801) was purchased from BD Transduction Laboratories^TM^, and the goat polyclonal anti-mouse cath-D (AF1029) from R&D Systems. The anti-human cath-D antibodies M1G8 and D7E3 were previously described [19]. The mouse monoclonal anti-tubulin antibody (clone 236-10501, #A11126) was from Thermo Fisher Scientific, the mouse monoclonal anti-Myc (clone 9B11) from Ozyme, and the rabbit polyclonal anti-β actin antibody (#A2066) from Sigma-Aldrich. The horse anti-mouse immunoglobulin G (IgG)-horseradish peroxidase (#7076), and goat anti-rabbit IgG-HRP (#7074S) secondary antibodies were from Cell Signaling Technology. The donkey anti-goat HRP conjugated antibody (FT-1I7890) was from Interchim. The Alexa Fluor 488-conjugated anti-rabbit IgG (#Ab150077) was purchased from Abcam, and the Cy3-conjugated anti-mouse IgG (#SA00009.1) from Proteintech. Hoechst 33342 (#FP-BB1340) was from Interchim FluoProbes.

### Cell lines, cell lysis, ELISA, and western blotting

The MDA-MB-231 cell line was previously described [17]. The Hs578T, MDA-MB-453 and MDA-MB-468 breast cancer cell lines were obtained from SIRIC Montpellier Cancer. The SUM159 breast cancer cell line was obtained from Asterand (Bioscience, UK). HEK-293 cells were kindly provided by A. Maraver (IRCM, Montpellier), HMFs by J. Loncarek and J. Piette (CRCL Val d’Aurelle-Paul Lamarque, Montpellier, France) [20], and HUVECs by M. Villalba (IRMB, Montpellier). Cell lines were cultured in DMEM with 10% fetal calf serum (FCS; GibcoBRL) except the SUM159 cell line that was cultured in RPMI with 10% FCS. Primary murine breast cancer cells were generated from end-stage tumors of Cre^ERT2^, *Ctsd^fl/fl^;* MMTV-PyMT mice as described previously [71]. All animal procedures were approved by the legal authorities and ethics committee at the regional council of Freiburg (registration numbers G14/18 and G18/38) and were performed in accordance with the German law for animal welfare. PyMT cells were cultured in DMEM/F12 medium supplemented with 10% FCS, 2 mM L-glutamine, and 1% penicillin-streptomycin at 37 °C with 5% CO_2_. 3 µM 4-hydroxytamoxifen (OH-Tam, Sigma Aldrich) was added to induce Cre-mediated recombination in the mouse *Ctsd* gene resulting in a premature stop codon. Cell lysates were harvested in lysis buffer (50 mM HEPES [pH 7.5], 150 mM NaCl, 10% glycerol, 1% Triton X-100, 1.5 mM MgCl_2_, 1 mM EGTA) supplemented with cOmplete™ protease and phosphatase inhibitor cocktail (Roche, Switzerland) at 4°C for 20 min, and centrifuged at 13 000 x g at 4°C for 10 min. Protein concentration was determined using the DC protein assay (Bio-Rad). Cath-D was quantified in TNBC and PDX cytosols by sandwich ELISA, after coating with the D7E3 antibody (200 ng/well in PBS) and with the HRP-conjugated M1G8 antibody (1/80), and using recombinant cath-D (1.25-15 ng/ml), as previously described [12]. TNBC cytosols were previously prepared and frozen [72]. For western blotting, proteins were separated on 13.5% SDS PAGE and analyzed by immunoblotting.

### Secretome preparation

To prepare secretomes from MDA-MB-231/HMF co-cultures, cells (ratio of 1:5, respectively) were plated in 150 mm Petri dishes in DMEM with 10% FCS. At a 90% confluence, MDA-MB-231/HMF cells were washed extensively with phenol red- and serum-free medium to remove serum proteins and grown in the same medium buffered with 50 mM HEPES [pH 7.5] for 24h. After harvesting, protease inhibitors (1 mM EDTA, protease inhibitor cocktail (Complete; Roche Applied Science)) were immediately added to the 24h-conditioned medium that was then clarified by centrifugation (500 g for 5 min; 8,000 g for 30 min) and filtered (0.45 µM). The 24h-conditioned medium was then concentrated to 0.2 mg/ml through Amicon filters (3 kDa cut-off, Millipore), and incubated in cleavage buffer with or without pepstatin A (12.5 µM) at pH 5.5 and 37°C for 0 or 60 min. Samples were then concentrated again by 15% trichloroacetic acid/acetone precipitation.

### Mass spectrometry analysis of protein N-termini (TAILS) in cell culture samples

After incubation with or without pepstatin and precipitation, the four samples (60 µg of total protein per condition: with/without pepstatin A and 0/60-min incubation) were dissolved in 200 mM HEPES pH 8 and denatured at 65°C for 15 min in the presence of 2.5 M guanidinium chloride. Then, each sample was reduced with 10 mM Tris (2-carboxyethyl) phosphine, alkylated with 25 mM iodoacetamide, and labeled with one TMT label (126, 127N, 127C, 128N; TMT 10-plex kit 90110 from Thermo Scientific), dissolved in DMSO in a 1:5 (total protein/ TMT label) mass ratio for 60 min. Labeling reactions were stopped by incubation with 5% hydroxylamine (Sigma) for 30 min, and the four samples were mixed and precipitated with cold methanol/acetone (8:1) (v/v). After two washes with cold methanol, the pellet was resuspended in 100 mM HEPES at pH 8 at a final protein concentration of 2 mg/ml and digested with trypsin (trypsin/total protein (1:100); Trypsin V511A, Promega) overnight. N-terminal peptide enrichment was performed on the digested sample by removing the internal tryptic peptides with a 1:5 mass excess of dialyzed HPG-ALD polymer (Flintbox, University of British Columbia). Enriched N-terminal peptides were then desalted with a C18 spin column (Thermo Fisher Scientific) and the eluate fraction was freeze-dried, resuspended in 0.1% formic acid and analyzed by LC-MS/MS on a Q-Exactive HF mass spectrometer (three replicates), as described for ATOMS experiments. Data files were analyzed with Proteome Discover 2.4 using the SEQUEST HT algorithm against the human protein database (SwissProt release 2019-12, 43835 entries). Precursor mass tolerance and fragment mass tolerance were set at 10 ppm and 0.02 Da, respectively, and up to 2 missed cleavages were allowed. Oxidation (M, P), pyroglutamate N-term (Q, E), acetylation (Protein N-terminus), and TMT6Plex (N-term, K) were set as variable modifications, and carbamidomethylation (C) as fixed modification. The without/with pepstatin A ratios were calculated for the two time points (0 min (126/127N) and 60 min (127C/128N)) and the ratios at 60 min were normalized to the ratios at 0 min. Only peptides with N-terminal TMT labeling and ratios showing at least a two-fold change, corresponding to more than 3-fold the standard deviation of the normal distribution of natural N-termini (Supplementary Fig. 1), were considered to indicate high confidence cleavage sites (Table 1 and Supplementary Table 1). The mass spectrometry proteomics data have been deposited in ProteomeXchange Consortium via the PRIDE [73] partner repository with the dataset identifier PXD022826 and 10.6019/PXD022826. Reviewer account details for data access: Username: reviewer; Password: wGObfZvw.

### SPARC cleavage by cath-D *in vitro* and *in cellulo*

Recombinant 52-kDa pro-cath-D (4 µM; R&D Systems) was auto-activated to 51-kDa pseudo-cath-D in 0.1 M Na-acetate buffer (pH 3.5), 0.2 M NaCl at 37°C for 15 min, as previously described [55]. Recombinant SPARC (1 µM; R&D Systems) was incubated with self-activated pseudo-cath-D (5 nM) at 37°C at different pH values in cleavage buffer [34 mM Britton-Robinson buffer in the presence of 0.12 mM phosphatidylcholine (Sigma-Aldrich) and 0.05 mM cardiolipin (Sigma-Aldrich) with or without 2 µM pepstatin A (Sigma-Aldrich)]. Cleaved SPARC peptides were separated by 13.5% or 17% SDS PAGE and analyzed by immunoblotting or silver staining (GE Healthcare Life Sciences), respectively. For *in cellulo* SPARC cleavage, 200, 000 MDA-MB-231 cells were plated with 100 000 HMFs in T25 cell culture flasks. After 24 h, culture medium was changed. Conditioned medium from co-cultured MDA-MB-231 cells and HMFs was obtained by adding DMEM without sodium bicarbonate and buffered with 50 mM HEPES buffer (pH 7.5) and without FCS for 24h. The 24h conditioned medium was then incubated, with or without pepstatin A (12.5 µM), at 37°C in cleavage buffer. Then, proteins in the medium (40 µl) were separated by 13.5% SDS-PAGE and analyzed by immunoblotting. In other *in cellulo* SPARC cleavage experiments, 200, 000 Hs578T, SUM159 or PyMT cells were incubated in DMEM without sodium bicarbonate and buffered with 50 mM HEPES buffer [pH 7.5] and without FCS for 24h, and the conditioned medium was analyzed as described above.

### Identification of cath-D-generated fragments by ATOMS

Recombinant SPARC (4 µM, 6 µg) was incubated with auto-activated 51-kDa or mature 34+14-kDa cath-D (200 nM) in 100 mM Na-acetate buffer (pH 5.9)/0.2 M NaCl with or without pepstatin A (200 µM; Sigma-Aldrich) in the presence of phosphatidylcholine (0.12 mM; Sigma-Aldrich) and cardiolipin (0.05 mM; Sigma-Aldrich) at 37°C for 15 min. SPARC cleavage was analyzed by 13.5% SDS PAGE and silver staining (GE Healthcare Life Sciences). The corresponding samples with/without pepstatin A (5 µg) were then processed for iTRAQ-ATOMS, as previously described [74]. Briefly, samples were denatured in 2.5 M guanidine hydrochloride and 0.25 M HEPES pH 8.0 at 65 °C for 15 min, reduced with 1 mM TCEP at 65 °C for 45 min, and alkylated with iodoacetamide at room temperature in the dark for 30 min. After iTRAQ labeling in DMSO, the two samples with/without pepstatin A were mixed and precipitated with eight volumes of freezer-cold acetone and one volume of freezer-cold methanol. The pellet was washed extensively with cold methanol, dried and resuspended in 5 µl of 50 mM NaOH. The pH was adjusted to 8 with 1.8 M HEPES pH 8.0, and the sample was digested at 37 °C with sequencing-grade trypsin (Promega; 1:50 protease:protein w/w ratio) or at 25°C with Glu-C (Promega; 1:20 protease:protein w/w ratio) overnight. After desalting on a C18 column (Pierce), the sample was analyzed by LC-MS/MS on a Q-Exactive HF mass spectrometer operated with the Xcalibur software (version 4.0) and equipped with a RSLC Ultimate 3000 nanoLC system (Thermo Scientific), as previously described [75]. Data files were analyzed with Proteome Discover 1.4 using the MASCOT (2.2 version) algorithm against the human protein database (SwissProt release 2017-01, 40500 entries including reverse decoy database). Precursor mass tolerance was set at 10 ppm and fragment mass tolerance was set at 0.02 Da, and up to 2 missed cleavages were allowed. Oxidation (M), deamidation (NQ), acetylation (Protein N-terminus), and iTRAQ 8Plex (N-term, K) were set as variable modifications, and carbamidomethylation (C) as fixed modification. Peptides and proteins were filtered using Percolator and a false discovery rate (FDR) of 1%. Peptides with N-terminal iTRAQ labeling were manually validated. Quantification was performed with the Reporter Ions Quantifier node. The peak integration was set to the Most Confidence Centroid with 20 ppm Integration Mass Tolerance on the reporter ions. The cath-D without pepstatin A/cath-D with pepstatin A ratios were calculated and ratios showing at least a two-fold change are conserved in Table 1 except for peptides corresponding to the mature N-terminus.

### Study approval

For TMA, TNBC samples were provided by the biological resource center (Biobank number BB-0033-00059) after approval by the Montpellier Cancer Institute Institutional Review Board, following the Ethics and Legal national French regulations for patient information and consent. For TNBC cytosols, patient samples were processed according to the French Public Health Code (law n°2004-800, articles L. 1243-4 and R. 1243-61). The biological resources center has been authorized (authorization number: AC-2008-700; Val d’Aurelle, ICM, Montpellier) to deliver human samples for scientific research. All patients were informed before surgery that their surgical specimens might be used for research purposes. The study approval for PDXs was previously published [41].

### Construction of tissue microarrays

Tumor tissue blocks with enough material at gross inspection were selected from the Biological Resource Centre. After hematoxylin-eosin-safranin (HES) staining, the presence of tumor tissue in sections was evaluated by a pathologist. Two representative tumor areas, to be used for TMA construction, were identified on each slide. A manual arraying instrument (Manual Tissue Arrayer 1, Beecher Instruments, Sun Prairie, WI, USA) was used to extract two malignant cores (1 mm in diameter) from the two selected areas. When possible, normal breast epithelium was also sampled as internal control. After arraying completion, 4 µm sections were cut from the TMA blocks. One section was stained with HES and the others were used for IHC.

### TMA immunohistochemistry

For SPARC and cath-D immunostaining, serial tumor sections from a TNBC TMA were incubated with 0.2 µg/ml anti-human SPARC mouse monoclonal antibody (clone AON-5031) for 30 min or with 0.4 µg/ml anti-human cath-D mouse monoclonal antibody (clone C-5) for 20 min after heat-induced antigen retrieval with the PTLink pre-treatment (Dako) and the High pH Buffer (Dako) and endogenous peroxidase quenching with Flex Peroxidase Block (Dako). After two rinses in EnVision^TM^ Flex Wash buffer (Dako), sections were incubated with a HRP-labeled polymer coupled to a secondary anti-mouse antibody (Flex® system, Dako) for 20 min, followed by incubation with 3,3’-diaminobenzidine as chromogen. Sections were counterstained with Flex Hematoxylin (Dako) and mounted after dehydration. Sections were analyzed independently by two experienced pathologists, both blinded to the tumor characteristics and patient outcomes at the time of scoring. SPARC signal was scored as low (<50%), or high (>50%), and cath-D signal was scored as low (<50%), or high (>50%) in cancer and stromal cells.

### Fluorescence microscopy

Paraffin-embedded PDX1995 tissue sections were deparaffined, rehydrated, rinsed, and saturated in PBS with 5% FCS at 4°C overnight. Sections were co-incubated with 1.2 µg/ml anti-SPARC rabbit polyclonal antibody (15274-1-AP) and 0.4 µg/ml anti-cath-D mouse monoclonal antibody (clone C-5) followed by co-incubation with AlexaFluor 488-conjugated anti-rabbit IgG (1/400) and a Cy3-conjugated anti-mouse IgG (1/500). Nuclei were stained with 0.5 µg/ml Hoechst 33342. Sections were then imaged with a 63X Plan-Apochromat objective on z stacks with a Zeiss Axio Imager light microscope equipped with Apotome to eliminate out-of-focus fluorescence. For co-staining of SPARC/cath-D, series of three optical sections (0.25 µm thick) were collected and projected onto a single plane.

### siRNA transfection

The siRNA duplex (21 nucleotides) against human cath-D siRNA (ID 4180) was purchased from Ambion (Austin, TX), and the firefly luciferase (Luc) siRNA (target sequence AACGUACGCGGAAUACUUCGA) was synthesized by MWG Biotech S.A [76]. MDA-MB-231 cells in 6-well plates were transiently transfected with 4 µg of siRNA using Lipofectamine 2000 (Invitrogen). At 48h post-transfection, 200 000 siRNA-transfected MDA-MB-231 cells were plated with 100 000 HMFs in T25 cell culture flasks for co-culture experiments.

### RT-qPCR

For gene expression analysis, fresh tumor tissues were homogenized in an Ultra-Turrax instrument. RNA was isolated using the RNeasy Mini Kit (Qiagen, Hilden, Germany) and 1 µg of total RNA was reverse transcribed using the iScript™ cDNA Synthesis Kit (Bio-Rad, Feldkirchen, Germany). Real-time PCR was performed using Platinum SYBR Green qPCR Super Mix-UDG (Life Technologies, Darmstadt, Germany) on a CFX96 real-time PCR machine (Bio-Rad) with the following primers: *SPARC* forward: 5’-GCTGTGTTGGAAACGGAGTTG-3’; *SPARC* reverse: 5’-CTTGCCATGTGGGTTCTGACT-3’; *ACTB* (β-actin) forward: 5’-ACCCAGGCATTGCTGACAGG-3’, *ACTB* reverse: 5’-GGACAGTGAGGCCAGGATGG-3’. *SPARC* expression data were normalized to *ACTB* expression.

### Expression and purification of recombinant proteins

The cDNA encoding human SPARC (303 amino acids according to the GenBank reference NP_003109) and its truncated fragments were PCR-amplified using the pcDNA3.1-SPARC plasmid as template [77], cloned into pGEM®-T Easy Vector (Promega), and then into the pSec-Tag2/hygroA vector (Thermo Fisher Scientific) by *Not* I digestion. Orientation and sequence were verified (Supplementary Table 2). Human embryonic kidney 293 (HEK-293T) cells were stably transfected with the plasmids using Lipofectamine 2000 (Invitrogen) according to the manufacturer’s instructions, and were selected with 400 µg/ml hygromycin B Gold^TM^ (Invivogen). Recombinant His-tagged proteins were purified from cell lysates on a nickel-chelating column (Ni-nitrilotriacetic acid agarose; His-select high flow nickel affinity gel; Sigma-Aldrich), as described previously [53]. The isolated recombinant proteins were analyzed by western blotting using anti-mouse Myc (clone 9B11) and anti-SPARC (clone AON-5031) antibodies and quantified using the Image J densitometric software (National Institutes of Health). To immunodeplete purified SPARC or its fragments, protein supernatants were incubated with an anti-Myc antibody (clone 9B11) overnight and protein G-Sepharose at 4°C for 4h, and supernatants were analyzed by immunoblotting to validate SPARC depletion. SPARC-immunodepleted supernatants were used as internal controls in the biological assays.

### Cell adhesion, migration, endothelial transmigration and invasion assays

Adhesion of MDA-MB-231 cells was assessed as described [53]. Briefly, 96-well plates were coated with fibronectin (10 µg/ml; sc-29011; Santa Cruz Biotechnology) at 4°C overnight, and saturated with 1% BSA in PBS. MDA-MB-231 cells were detached with HyQTase (HyClone), washed in DMEM without FCS, and 1.5 10^5^ cells were pre-incubated or not with SPARC or its cleaved fragments at room temperature for 10 min. Cells (5 10^4^ cells) were plated and left in serum-free medium at 37°C for 30 min. Non-adherent cells were removed by floatation on a dense Percoll solution containing 3.33% NaCl (1.10 g/l), and adherent cells were fixed (10% [vol/vol] glutaraldehyde) using the buoyancy method [78]. Cells were stained with 0.1% crystal violet, and absorbance was measured at 570 nm. For migration assays, 8-µm pore Transwell inserts in 24-well plates (polyvinyl pyrrolidone-free polycarbonate filter) (Corning Inc., Corning, NY, USA) were coated with 10 µg/ml fibronectin (500 ng) at 4°C for 24h. For invasion assays, 8-µm pore Transwell inserts were coated with Matrigel (100 µg, Corning). MDA-MB-231 cells (2 10^5^ cells) were pre-incubated or not with SPARC or its cleaved fragments at room temperature for 10 min, and then plated (5 10^4^ cells/well) in FCS-free DMEM on the coated insert in the upper chamber. For transmigration assay, 10^5^ HUVECs were plated in the upper chamber of a gelatin-coated Transwell insert and grown in complete endothelial medium to confluence, as previously described [26]. The endothelial monolayer was then incubated with human TNFα (10 ng/ml; PeproTech) for 16h. MDA-MB-231 cells (3 10^5^ cells), pre-incubated or not with SPARC or its cleaved fragments at room temperature for 10 min, were then plated (10^5^ cells/well) in FCS-free DMEM on top of the endothelial monolayer. In these different assays, DMEM supplemented with 10% FCS was used as chemoattractant in the bottom chamber. After 16h, non-migrating/non-invading/non-transmigrating cells on the apical side of each insert were scraped off with a cotton swab, and migration, invasion and transmigration were analyzed with two methods: (1) migrating/invading/transmigrating cells were fixed in methanol, stained with 0.1% crystal violet for 30 min, rinsed in water, and imaged with an optical microscope. Two images of the pre-set field per insert were captured (x100); (2) migrating/invading/transmigrating cells were incubated with 3-(4,5-dimethylthiazol-2-yl)-2,5-diphenyltetrazolium bromide (MTT; 5 mg/ml, 1/10 volume; Sigma-Aldrich) added to the culture medium at 37°C for 4h. Then, the culture medium/MTT solution was removed and centrifuged at 10 000 rpm for 5 min. After centrifugation, cell pellets were suspended in DMSO. Concomitantly, 300 µl of DMSO was added to each well and thoroughly mixed for 5 min. The optical density values of stained cells (cell pellet and corresponding well) were measured using a microplate reader at 570 nm.

## ACKNOWLEDGMENTS

This work was supported by a public grant overseen by the French National Research Agency (ANR) as part of the “Investissements d’Avenir” program (reference: LabEx MabImprove ANR-10-LABX-53-01), SIRIC Montpellier Cancer Grant INCa_Inserm_DGOS_12553, University of Montpellier, the associations ‘Ligue Régionale du Gard’, ‘Ligue Régionale de l’Herault’, ‘Ligue Régionale de la Charente Maritime, and ‘Association pour la Recherche sur le Cancer’ (ARC). The work of TR was supported by the German Research Foundation (DFG) SFB 850 project B7, RE 1584/6-2, and GRK 2606. We thank Susanne Dollwet-Mack and Lisa Heß (Institute of Molecular Medicine and Cell Research, Freiburg, Germany) for helpful discussions and support. The authors acknowledge Biological Resources Centre from Montpellier Cancer Institute (ICM Biobank n° BB 0033 00059).

## AUTHOR CONTRIBUTIONS

LBA, AM, DD, FD, CO, PR, TR, CM, ELC designed the experiments and prepared the manuscript. LBA, AM, TD, DD, FD, CD, FB, JSF, PFH, PR, TR, ELC performed the experiments. LBA, AM, DD, FD, JSF, SDM, PFH, CO, STD, WJ, SG, PR, TR, CM, ELC provided material and analyzed data. LBA, AM, TC, TR, CM, ELC analyzed data and proof-read and finalized the manuscript.

