## supplemental text for "A 9-kDa matricellular SPARC fragment released by cathepsin D exhibits pro-tumor activity in the triple-negative breast cancer microenvironment"

### SUPPLEMENTARY MATERIALS AND METHODS

#### Gene expression data analysis

The 10-year recurrence-free survival was calculated using the on-line Kmplot tool accessed on October 2 2017 with the 200766_at Affymetrix probe ([40], www. http://kmplot.com). Analysis was restricted to the 255 patients with TNBC present in the database at this date and with the best cut-off option. The cut-off value for SPARC was 1890 with a probe range from 268 to 17781. The cut-off value for cath-D was 1919 with a probe range from 50 to 6518. Differences were evaluated with the log-rank test.

#### Fluorescence microscopy

MDA-MB-231 cells were fixed with 4% (vol/vol) paraformaldehyde at room temperature for 20 min, and permeabilized with 0.5% (vol/vol) Triton X-100 in PBS. Fixed cells were incubated with tetramethylrhodamine isothiocyanate-labeled phalloidin (Sigma-Aldrich) to visualize F-actin reorganization. DNA was stained with Hoechst 33342 (1/10 000). Cells were imaged with a 63X Plan-Apochromat objective on z stacks with a Zeiss Axio Imager light microscope equipped with Apotome to eliminate out-of-focus fluorescence.

### LEGEND TO SUPPLEMENTARY FIGURES AND TABLES

**Supplementary Fig. 1. Distribution of the normalized ratios for mature N-terminal peptides at 60 min.** 637 N-terminal peptides corresponding to the beginning of mature proteins (including after first methionine removal or signal peptide removal) were included because they are not supposed to be affected by cath-D and represent the read-out of the sample intrinsic variability (distribution standard deviation = 0.293). Dashed lines indicate the 2-fold changes in abundance ($\log 2 \leq -1$ or $\geq 1$) chosen as cut-offs to identify severely affected N-terminal regions.

**Supplementary Fig. 2. Kaplan-Meier curves of recurrence-free survival according to *CTSD* and *SPARC* mRNA expression in TNBC**

N=255 patients with TNBC; for *CTSD*, HR=1.65 [1.08-2.53], *P* = 0.019; for SPARC, HR=1.6 [0.91-2.79], *P* = 0.097, log-rank test. Probability, recurrence-free survival. Time, months after diagnosis.

**Supplementary Fig. 3. Cath-D and SPARC expression and secretion in an inducible *Ctsd* knock-out MMTV-PyMT mammary tumor cell line**

Cells were incubated or not with 4-hydroxytamoxifen (OH-Tam, 3 μM) for 4 days to induce *Ctsd* knock-out. Culture medium was changed and cells were incubated in the absence of serum for 24h. Whole cell extracts (left panel) and 24-hour conditioned media (right panel) were separated by 13.5% SDS-PAGE and analyzed by immunoblotting with anti-SPARC (15274-1-AP) and anti-cath-D (AF1029) antibodies. β-actin, loading control.

**Supplementary Fig. 4. Analysis of cath-D-induced recombinant cleaved SPARC fragments**

Recombinant FL SPARC was incubated with recombinant auto-activated pseudo-cath-D at 37°C in cleavage buffer (at pH 5.5) with or without pepstatin A (Pepst) for 5h. After 5h, pepstatin A was added also in the samples without pepstatin A. FL SPARC and cath-D-induced cleaved SPARC fragments (80 ng) were analyzed by 17% SDS-PAGE and silver staining.

**Supplementary Fig. 5. FL SPARC effect on MDA-MB-231 cell adhesion is dose-dependent**

MDA-MB-231 cells were let to adhere on fibronectin in the absence (CTRL, PBS) or presence of increasing doses of recombinant FL SPARC (2, 5 or 10 μg/ml) for 30 min. Left panels, representative images of adherent cells. Right panel, adherent cells were stained with crystal violet, and adhesion was quantified by absorbance at 570 nm. Mean ± SD (n=3); ***, $p<0.001$, ANOVA and Bonferroni's post hoc test.

**Supplementary Fig. 6. FL SPARC effect on adhesion of cath-D-silenced TNBC cells**

**(A)** ***CSTD*** **silencing in MDA-MB-231 cells.** Cells were transfected with Luc or cath-D siRNAs. At 48h post-transfection, cellular (lysate) and secreted (CM, conditioned medium) cath-D in siRNA-transfected cells were analyzed by immunoblotting with two anti-cath-D antibodies (clone 49, #610801, and H-75, respectively).

**(B) FL SPARC effect on adhesion of siRNA-transfected MDA-MB-231 cells.** At 48h post-transfection, cath-D- or Luc-silenced cells were let to adhere on fibronectin in the absence or presence of recombinant FL SPARC at a final concentration of 240 nM for 30 min. Left panels, representative images of adherent cells. Right panel, adherent cells were stained with crystal violet, and adhesion was quantified by absorbance at 570 nm. CTRL, PBS. Mean ± SD; ns, not significant; ***, p<0.001, ANOVA and Bonferroni's post hoc test.

**(C) SPARC and cath-D in conditioned media of silenced-MDA-MB-231 cells.** After the adhesion assays, the conditioned media from cells transfected with Luc or cath-D siRNAs were analyzed with anti-cath-D (H-75) and anti-SPARC (15274-1-AP) antibodies. CTRL, PBS.

**Supplementary Fig. 7. Effects of FL SPARC and cath-D-induced cleaved SPARC fragments on TNBC cell spreading**

**(A) Cell spreading and F-actin distribution**. MDA-MB-231 cells were plated on fibronectin in the presence or not of recombinant FL SPARC, or recombinant cath-D-induced cleaved SPARC fragments (final concentration: 240 nM) for 30 min. Representative phase-contrast images of MDA-MB-231 cells (magnification x100) (top panels). F-actin was stained with phalloidin (red) and nuclei with 0.5 μg/ml Hoechst 33342 (blue) (middle and lower panels). Scale bar, 10 μm. Higher magnification of F-actin immunostaining (bottom panels). CTRL, PBS in cleavage buffer.

**(B) Quantification of spread cells**. Mean (% of spread cells) ± SD; *, p<0.05, ***, p<0.001, ANOVA and Bonferroni's post-test. Spread cells were quantified according to F-actin distribution.

**Supplementary Fig. 8. Structure of the 16-, 9- and 6-kDa C-terminal SPARC fragments**

**(A) Amino acid sequence of the 16-, 9- and 6-kDa C-terminal SPARC fragments in the extracellular $Ca^{2+}$ binding domain of SPARC**. The entire C-terminal extracellular $Ca^{2+}$ binding domain of human SPARC (amino acids 154-303) is shown. The sequence of the 9-kDa C-terminal SPARC fragment (amino acids 235-303) is in bold. The residues coordinating $Ca^{2+}$ ions, located in the two EF-hand motifs (EF-hand 1 amino acids 227-260; EF-hand 2 amino acids 262-294), are highlighted in yellow. Arrows, 16-, 9- and 6-kDa fragments.

**(B) Cartoon showing the localization of the 16-, 9- and 6-kDa C-terminal SPARC fragments within the FL SPARC protein.** The atomic coordinates of FL SPARC were found online on the Protein Data Bank (PDB) (DOI: 10.2210/pdb2V53/pdb). These images were generated using the Swiss PDB Viewer 4.0.4 software. The leading chains of the 16-, 9- and the 6-kDa SPARC fragments within FL SPARC are shown in pink, green, and blue, respectively. The residues coordinating $Ca^{2+}$ ions are in yellow.

**Supplementary Fig. 9. Analysis of recombinant FL SPARC, cath-D-induced recombinant SPARC fragments, and purified 9-kDa C-terminal SPARC fragment**

Left panel, recombinant FL SPARC (rec. SPARC) was incubated with auto-activated recombinant cath-D at 37°C in cleavage buffer (pH 5.5) with or without pepstatin A for 5h. After 5h, pepstatin A was added in the non-pepstatin A treated rec. SPARC sample. Immunoblot analysis using an anti-SPARC antibody (clone AON-5031) of purified FL SPARC incubated with cleavage buffer and pepstatin A, rec. FL SPARC, and cath-D-induced cleaved SPARC fragments incubated with SPARC immunodepleted supernatant from the 9-kDa SPARC fragment purification (240 nM each). Right panel, immunoblotting using an anti-Myc antibody (clone 9B11) of the levels of purified FL SPARC and 9-kDa C-terminal SPARC fragment in cleavage buffer and pepstatin A (240 nM each).

**Supplementary Fig. 10. Effects of FL SPARC, cath-D-induced cleaved SPARC fragments, and 9-kDa C-terminal SPARC fragment on TNBC cell spreading**

**(A) Cell spreading and F-actin distribution**. MDA-MB-231 cells were plated on fibronectin in the presence or not of recombinant FL SPARC, recombinant cath-D-induced cleaved SPARC fragments, or purified 9-kDa C-terminal SPARC fragment (final concentration: 240 nM) for 30 min. Representative phase-contrast images of MDA-MB-231 cells (magnification x100) (top panels). F-actin was stained with phalloidin (red) and nuclei with 0.5 µg/ml Hoechst 33342 (blue) (middle and bottom panels). Scale bar, 10 μm. Higher magnification of F-actin immunostaining (bottom panels). CTRL, PBS in cleavage buffer and SPARC immunodepleted supernatant from the 9-kDa SPARC fragment purification.

**(B) Spread cells**. Spread cells were quantified according to F-actin distribution and shown as the mean (% of spread cells) ± SD; *, $p<0.05$, ***, $p<0.001$, ANOVA and Bonferroni's post-test.

**Supplementary Table 1. Sequences of the fragments identified by TAILS**

**Supplementary Table 2. Primers for *SPARC* fragment cloning**
