## Supplementary material for "A 9-kDa matricellular SPARC fragment released by cathepsin D exhibits pro-tumor activity in the triple-negative breast cancer microenvironment": Supp Table 2

| Gene | Primer sequence (strand) | Product size (bp) |
|---|---|---|
| *Sparc 42-kDa* | 5'-AGCCCCTCAGCAAGAAGCC- 3' (+)<br>5'-GCGATCACAAGATCCTTGTC- 3' (-) | 884 |
| *Sparc 34-kDa* | 5'-AGCCCCTCAGCAAGAAGCC- 3' (+)<br>5'-GCAGCACGCAGTGGAGCCAG- 3' (-) | 720 |
| *Sparc 27-kDa* | 5'-AGCCCCTCAGCAAGAAGCC- 3' (+)<br>5'-GCCAGGCGCTTCTCATTC- 3' (-) | 567 |
| *Sparc 16-kDa* | 5'-ATATGAGAGGGATGAGGAC- 3' (+)<br>5'-GCGATCACAAGATCCTTG- 3' (-) | 369 |
| *Sparc 9-kDa* | 5'-ATTCGGCCAGCTGGACCAG- 3' (+)<br>5'-GCGATCACAAGATCCTTG- 3' (-) | 201 |
| *Sparc 6-kDa* | 5' -ACCCCTCATCCCCATGGAG- 3' (+)<br>5'-GCGATCACAAGATCCTTG- 3' (-) | 132 |

**Supplementary Table 2. Primers for *SPARC* fragment cloning**
